# Altered cleavage of Caspase-1 in hepatocytes limits control of malaria in the liver

**DOI:** 10.1101/2021.01.28.427517

**Authors:** Camila Marques-da-Silva, Barun Poudel, Rodrigo P. Baptista, Kristen Peissig, Lisa S. Hancox, Justine C. Shiau, Lecia L. Pewe, Melanie J. Shears, Thirumala-Devi Kanneganti, Photini Sinnis, Dennis E. Kyle, Prajwal Gurung, John T. Harty, Samarchith P. Kurup

## Abstract

Malaria, caused by *Plasmodium* parasites, is a devastating disease that kills over half a million people each year^1^. *Plasmodium* sporozoites inoculated by mosquitoes into mammalian hosts undergo a clinically silent phase of obligatory development and replication in hepatocytes before initiating the life-threatening blood-stage of malaria^2^. Thus, understanding the immune responses elicited by *Plasmodium* infection in the liver is key to controlling clinical malaria and transmission^3,4^. Here, we show that *Plasmodium* DNA can be detected by AIM2 (absent in melanoma 2) sensors in the infected hepatocytes, resulting in Caspase-1 activation and pyroptotic cell-death. However, Caspase-1 was observed to undergo only partial cleavage in hepatocytes, limiting pyroptosis, and the maturation of pro-inflammatory cytokines classically associated with Caspase-1 activation. We discovered that the extent of Caspase-1 cleavage in cells is determined by the expression of ASC (apoptosis-associated speck-like protein containing a CARD). ASC expression is inherently low in hepatocytes, and transgenically enhancing it in the hepatocytes induced complete processing of Caspase-1, efficient secretion of pro-inflammatory cytokines, enhanced pyroptotic cell-death, and markedly improved control of malaria infection in the liver. In addition to describing a novel pathway of natural immunity to malaria, our findings uncover a key aspect of liver biology that may have been exploited during evolution by successful hepatotropic pathogens.

## Main

*Plasmodium* is a complex eukaryotic pathogen that has evolved to flourish in two vertebrate host tissues uniquely replete with immune cells, the liver and the blood^2^. Although the immune responses elicited by *Plasmodium* infection in its mammalian hosts have been intensely studied, our understanding of the innate immune responses that control malaria infection in the liver is still rudimentary^5^. Considering that the development of *Plasmodium* in the liver is a prerequisite for clinical malaria and transmission, and the key target of frontline anti-malarial vaccines, understanding of the immune responses generated against *Plasmodium* in the liver is critical to combating malaria^3,4,6^.

### AIM2 inflammasome controls malaria in the liver

To identify immune pathways elicited by *Plasmodium* in hepatocytes in an unbiased manner, we compiled the transcriptional signature of human hepatocytes infected with *P. falciparum* (*Pf*) through single-cell RNA sequencing (Extended Data Fig 1a-b). Expression of genes encoding molecules involved in major biochemical pathways including host responses to cell-injury, cell-adhesion, acute phase responses and inflammation were altered in *Pf* infected hepatocytes (Extended Data Fig 1c-f). Although various genes associated with cell-damage, programmed celldeath and inflammatory pathways were transcriptionally altered (Extended Data Fig 1g) few were more uniformly and profoundly impacted as the genes of the inflammasome pathway (Fig 1a). Inflammasome activation in host cells is integral to the control of intracellular pathogens^7^. Supporting the transcriptional data, formation of inflammasome complexes with Caspase-1 was observed in primary human and mouse hepatocytes infected with *Pf* or *P. yoelii* (*Py*) respectively (Fig 1b-c). Caspase-1 was found to associate with *Plasmodium* early in their development in the hepatocytes (Extended Data Fig 2), and its activation was mediated through the ASC adaptor (Extended Data Fig 3b). ASC or Caspase-1 deficiency resulted in higher *Py* burden in the liver (Fig 1d) indicating that the inflammasome pathway makes a substantial contribution to parasite control in the liver. Of note, mice deficient in Caspase-1 (Casp1KO) used in this study are also deficient in Caspase-11^8^. However, Caspase-11 was not activated by *Py* infection in hepatocytes, nor did it impact the control of *Py* infection in the liver (Extended Data Fig 4). Therefore, these data indicated that Caspase-1 activation was critical for protection from malaria in the liver.

**Figure 1:**
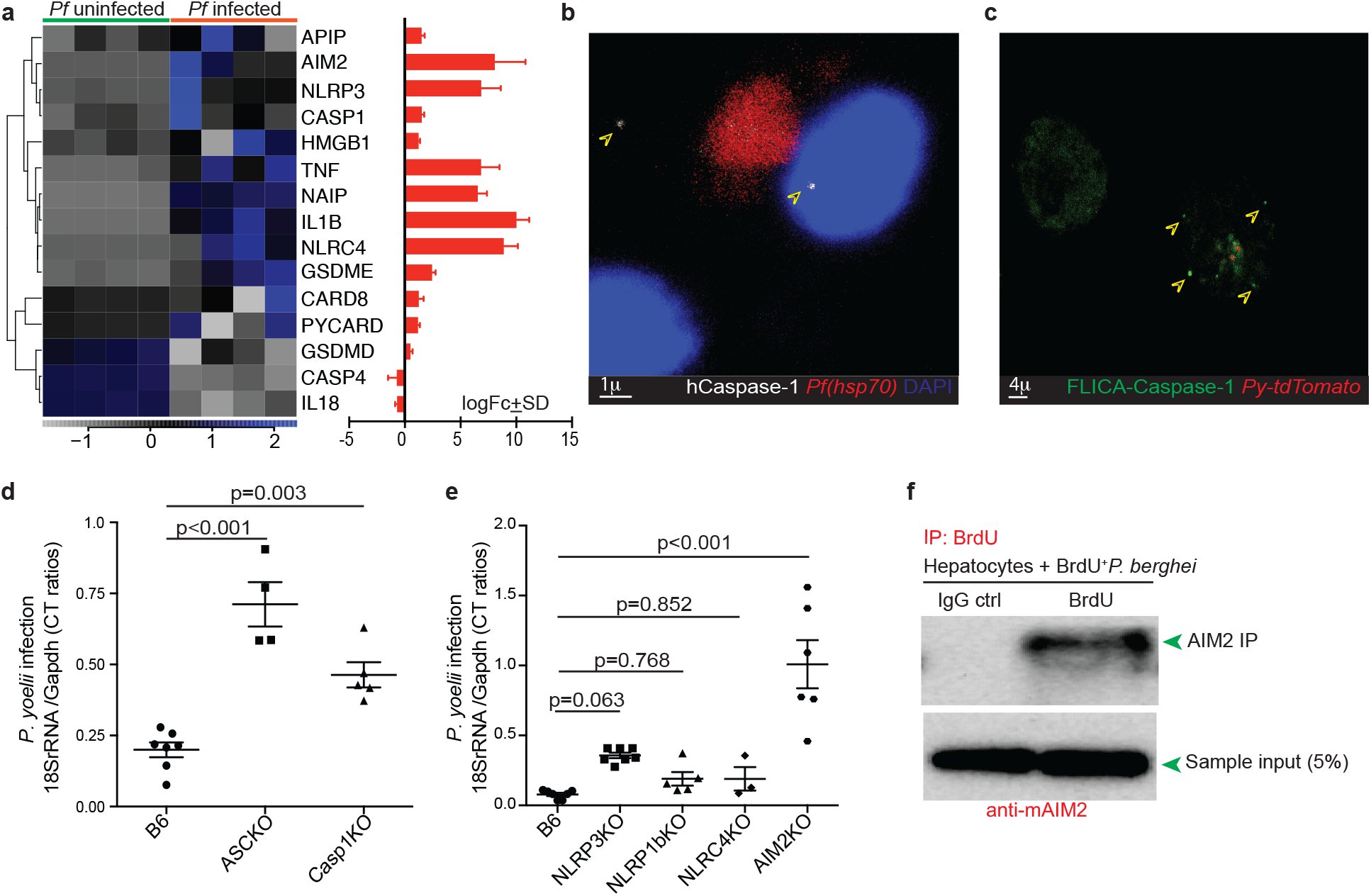
AIM2 mediated inflammasome activation in hepatocytes control liver-stage malaria. **a**, Heat map depicting transcriptional differences in the genes of the canonical inflammasome pathway in primary human hepatocytes, maintained ex vivo and infected with *Pf*, 4d p.i. The bar graph on right shows log fold-change + standard deviation (logFC + SD) in the differential expression of the indicated genes. **b**, Representative pseudo-colored confocal image depicting Caspase-1 activation (arrows) in human hepatocytes in a section of the liver of a humanized mouse inoculated with *Pf* (30h p.i) and stained with the indicated antibodies. **c**, Representative pseudo-colored confocal image depicting Caspase-1 activation (FLICA^+^, arrows) in ex vivo cultured primary mouse hepatocytes infected with *Py* (expressing tdTomato), 30h p.i. **d-e,** Scatter plots showing relative liver-parasite burdens in the indicated mice inoculated with *Py*, at 36h p.i. Dots represent individual mice. Data presented as mean + s.e.m and analyzed using ANOVA with Dunnett’s corrections, yielding the indicated p values. **e**, Immunoblot analysis for AIM2 after immunoprecipitation of the whole-cell lysates of primary mouse hepatocytes infected with BrdU^+^ *Pb* (24h p.i.), using with anti-BrdU antibodies. Data in b-e represent >3 separate experiments.

The dependence of protection on ASC suggested that *Plasmodium* in hepatocytes may be sensed directly by NLRP3, NLRP1b, NLRC4, or AIM2 pattern recognition receptors (PRRs)^7^. To determine which of these sensors were pertinent to the control of *Plasmodium* in the liver, we compared *Py* infections in the livers of mice genetically deficient for NLRP3, NLRP1b, NLRC4, or AIM2, and control B6 mice. Only AIM2KO mice exhibited higher *Py* burden in the liver (Fig 1e). *Py* infection itself enhanced the expression of AIM2 in hepatocytes in a type I interferon (IFN) dependent manner (Extended Data Fig 5a). Type 1 IFNs induced in hepatocytes consequent to *Plasmodium* infection are known to promote control of liver-stage malaria^9^. Our observation that type 1 IFNs induce the expression of AIM2 in hepatocytes offers a potential pathway for type 1 IFN driven immunity to liver-stage malaria. Of note, type 1 IFN treatment also was sufficient to induce AIM2 expression in bone marrow derived macrophages (BMDMs) (Extended Data Fig 5b). Though hepatocytes are the only cells known to support *Plasmodium* replication and development in the liver, CD11c^+^ dendritic cells (DCs) recruited to the liver following *Plasmodium* infection can acquire *Plasmodium* parasites from previously infected hepatocytes^10^. However, Caspase-1 or AIM2 in hematopoietic cells did not contribute to the control of *Plasmodium* infection in the liver, as indicated by *Py* infection in reciprocal bone-marrow chimeric mice generated from Casp1KO, AIM2KO and B6 mice (Extended Data Fig 6).

The only known ligand for AIM2 is double-stranded (ds) DNA^11^. Hence, we surmised that *Plasmodium* derived dsDNA is directly sensed by AIM2 receptors in hepatocytes. To query this possibility, we determined if AIM2 could be co-immunoprecipitated with BrdU^+^ cytosolic DNA to demonstrate direct association between AIM2 and parasite DNA in cells. Indeed, anti-BrdU mediated pulldown in primary hepatocytes transfected with BrdU^+^ genomic DNA coimmunoprecipitated AIM2 (Extended Data Fig 7a). We then examined the possible direct association of *Plasmodium* DNA with hepatocyte AIM2 by incorporating BrdU into the DNA of the sporozoite stage of *P. berghei* (*Pb*). BrdU is known to integrate into the DNA of *Plasmodium* parasites cultured in media replete with BrdU^12,13^. *Pb* infected *Anopheles* mosquitoes were reared on BrdU laced sugar-water. As a result, high frequencies (94+2%) of BrdU^+^ *Pb* sporozoites that could give rise to liver-stages of *Pb* with BrdU incorporated DNA were generated (Extended Data Fig 7b-d), and used to infect primary hepatocytes. Co-immunoprecipitation of BrdU and AIM2 indicated that *Plasmodium* DNA directly associated with hepatocyte AIM2 sensors in infected cells (Fig. 1f). Taken together, these data indicate that the AIM2 sensors in hepatocytes facilitate detection and control *Plasmodium* infection in the liver through inflammasome mediated Caspase-1 activation.

### Gasdermin D effects *Plasmodium* control in liver

Broadly, Caspase-1 activation has two discrete immunological functions in host cells: (1) induction of programmed cell-death through proteolytic activation of Gasdermin D (GSDMD) to generate membrane pores and (2) concurrent proteolytic maturation and release of the proinflammatory cytokines, IL-1 and IL-18 through these pores^14^. *Py* infection induced detectable cell-death in hepatocytes, mediated by AIM2 and GSDMD (Fig 2a). The characteristic vacuolation and rupture of hepatocytes following *Py* infection in a GSDMD dependent manner suggested that *Py* infection induced pyroptotic cell-death in infected hepatocytes (Fig 2b, Supplementary videos S1, S2)^15^. *Py* infection induced GSDMD activation in hepatocytes in a Caspase-1 dependent manner (Fig 2c), with GSDMD localization suggesting the formation of plasma membrane pores (Fig 2d). Consistent with these findings, genetic deficiency, or therapeutic inhibition of GSDMD with disulfiram^16^ resulted in suboptimal control of malaria infection in the liver (Fig. 2e-f). Taken together, these data showed that the AIM2-Caspase-1 axis induced GSDMD-mediated pyroptotic cell-death in *Plasmodium* infected hepatocytes, aiding in the control of liver-stage malaria. Rather surprisingly however, Caspase-1 activation in hepatocytes induced reduced levels of mature IL-1β or IL-18 (Extended Data Fig 8a-b). Maintaining the capacity to use the inflammasome pathway to eliminate intracellular pathogens through cell-death in hepatocytes, while concurrently producing less pro-inflammatory cytokines may be perceived as an evolutionary adaptation to defend the liver from pathogenic insults without sacrificing the liver’s immunotolerant nature.

**Figure 2:**
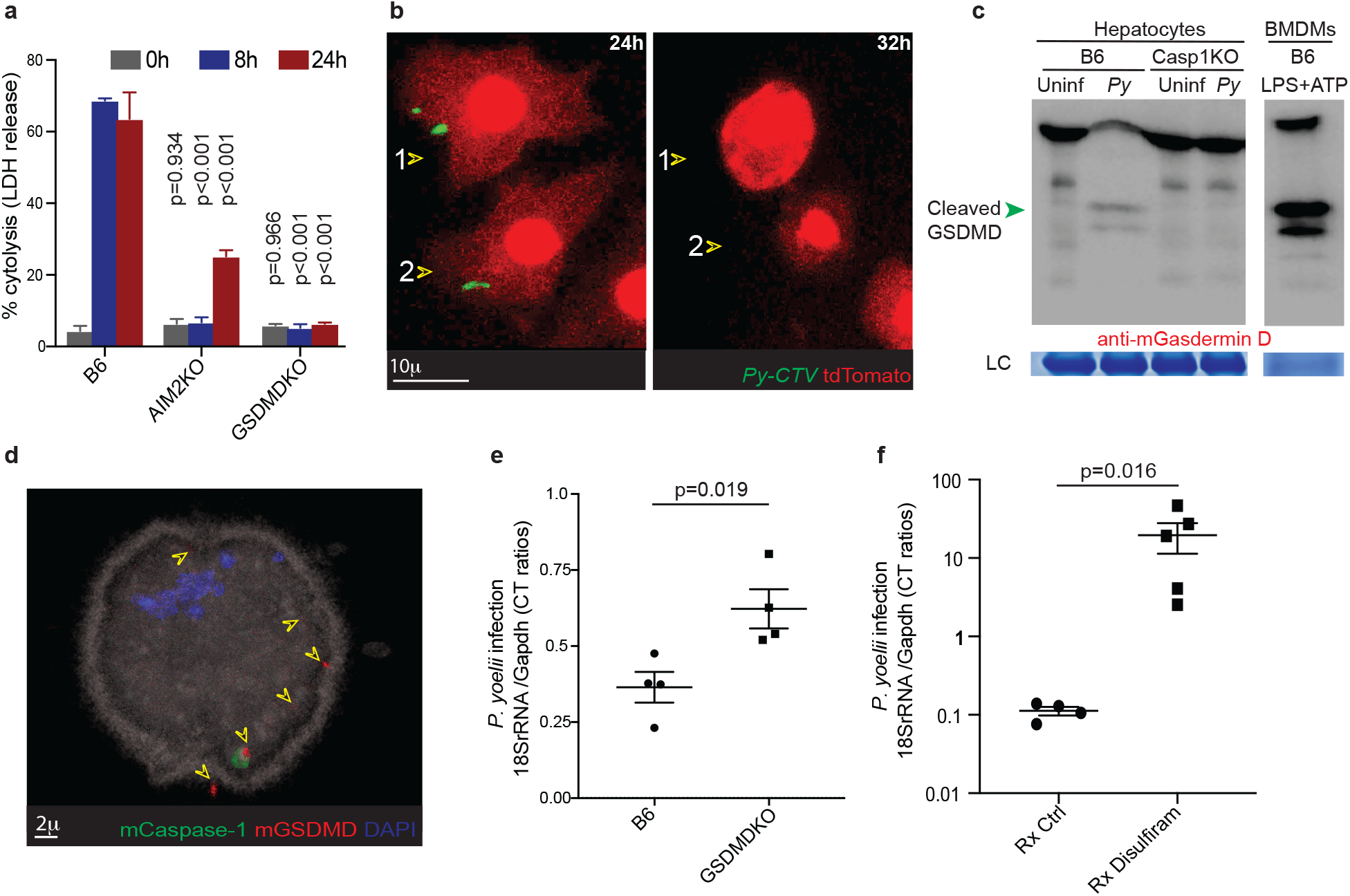
Gasdermin D mediated cell-death in *Plasmodium* infected hepatocytes. **a,** Comparison of cytolysis determined by LDH release assay in ex vivo cultured primary hepatocytes derived from the indicated mice, co-incubated with *Py* for the indicated times. Data presented as mean + s.e.m, analyzed using ANOVA with Dunnett’s correction comparing each time point to the corresponding one in B6 mice, to yield the presented p-values. **b**, Representative confocal time-lapse images showing the same primary tdTomato^+^ B6 mouse hepatocytes infected with *Py* (CellTrace Violet^+^, arrows) observed at 24h and 32h of co-incubation. Hepatocyte #1 and #2 shown on left just prior to undergoing pyroptotic rupture; see Movie S1 for complete sequence of events. **c**, Immunoblot analysis for GSDMD cleavage in primary hepatocytes obtained from the indicated mice co-incubated with *Py*, at 24h. BMDMs treated with LPS+ATP (4h) served as the positive control. LC: loading control. **d**, Representative pseudo-colored high-resolution confocal image of a B6 mouse hepatocyte co-cultured with *Py* for 30h showing nuclear disintegration indicating cell-death and GSDMD localization (arrows) to the plasma membrane. **e**, Scatter plots showing relative liver-parasite burdens in the indicated mice inoculated with *Py*, 36h p.i. **f**, Scatter plots showing relative liver-parasite burdens in control or Disulfiram treated (−1d, 0, 1d p.i) mice inoculated with *Py*, determined at 42h p.i. **e-f**, Data presented as mean + s.e.m, analyzed with 2-tailed t-tests to yield the presented p value. All data shown represent >3 separate experiments.

### Incomplete activation of Caspase-1 in hepatocytes

Activation of Caspase-1 occurs through the autoproteolytic cleavage of its tripartite progenitor procaspase-1 at the macromolecular inflammasome complex assembled in the cytosol following PRR stimulation^7^. In the current model for Caspase-1 activation developed based on studies in immune cells or *in vitro* cultured cell-lines, ASC forms the foundation of such inflammasome complexes (ASC specks), where constitutively expressed procaspase-1 molecules congregate to undergo self-cleavage^7,17–19^. Cleavage of the 46kDa procaspase-1 (p46) in immune cells or cell lines generate separate CARD, p20 (20kDa) and p10 (10kDa) domains (Extended Data Fig. 9a). Subsequent to this, catalytically functional hetero-tetramers composed of p20 and p10 subunits are generated, that carry out the downstream biological functions of Caspase-1^11,20^. In the case of macrophages cultured *in vitro*, a short-lived (<30 mins) CARD-p20 intermediate product has also been reported^17^. In stark contrast to this existing paradigm, we observed that Caspase-1 activation in mouse or human primary hepatocytes infected with *Plasmodium*, or stimulated with LPS and ATP (the standard pathogen associated molecular patterns used in studies in immune cells or cell lines) produced a novel 32 kDa (p32) Caspase-1 cleavage product, without detectable p10 or p20 (Fig. 3a-c). To test if p32 Caspase-1 is an artifact of *in vitro* culture or stimulation, we infected liver-humanized chimeric mice with *Pf* sporozoites, in which, the *Pf* infection would be limited to the hepatocytes of human origin^21^. p32 was the only cleaved form of Caspase-1 immunoprecipitated from the whole liver lysates of these mice (Fig 3d). Unlike in BMDMs stimulated with LPS+ATP, where distinct p20 and p10 cleavage products were detected, *Py* infected hepatocytes presented only p32 when probed with either p20 or p10 Caspase-1 subunit-specific antibodies (Fig. 3e). This suggested that the p32 observed in hepatocytes may be composed of unseparated p20 and p10 domains of Caspase-1 (Extended Data Fig 9a). To objectively test this possibility, we employed p20 subunit specific antibody to immunoprecipitate Caspase-1 from the lysates of *Py* infected hepatocytes, resolved it on a denaturing gel and probed with p10 subunit specific antibody (Extended Data Fig. 9b). The observation of p32 in this experiment confirmed that p32 was indeed composed of both p20 and p10 domains, possibly linked by an uncleaved interdomain linker (IDL, Extended Data Fig. 9a). Although various groups have examined Caspase-1 activation in the liver, the majority of these studies were conducted using whole-liver extracts containing myeloid as well as hepatic cells^22,23^, or cell-lines where celldeath pathways are often dysregulated^24–27^. In those instances where primary hepatocytes were discretely examined in the context of infections or aseptic inflammatory diseases of the liver, conventional cleavage of Caspase-1 was found to be limited to the non-parenchymal cells alone^28–30^. The novel p32 Caspase-1 cleavage product generated in *Py* infected primary hepatocytes remained detectable for over 24h post infection, was ASC-dependent, and catalytically functional (Extended Data Fig. 9c-d). These observations clearly distinguished the p32 Caspase-1 in hepatocytes from the short-lived, 33kDa CARD-p20 intermediate generated during Caspase-1 processing in macrophages^17^. Infection with *Salmonella enterica* Typhimurium (*St*) or stimulations with LPS and Nigericin also generated the p32 cleavage product in hepatocytes, unlike such treatments in BMDMs (Fig. 3f), reiterating that the incomplete processing of Caspase-1 is a characteristic of hepatocytes, rather than of *Plasmodium* infection itself. Delivery of *Py* DNA into primary hepatocytes by transfection induced p32 Caspase-1 in an AIM2 dependent manner, corroborating our previous finding that *Plasmodium* DNA induces Caspase-1 activation through AIM2 mediated signaling (Extended Data Fig. 10). Our findings challenged the universality of the existing model of Caspase-1 processing in cells and revealed that pyroptosis and secretion of inflammatory cytokines are not inseparable functional consequences of inflammasome formation and Caspase-1 activation. However, the underlying mechanism behind this functional dichotomy remained unknown.

**Figure 3:**
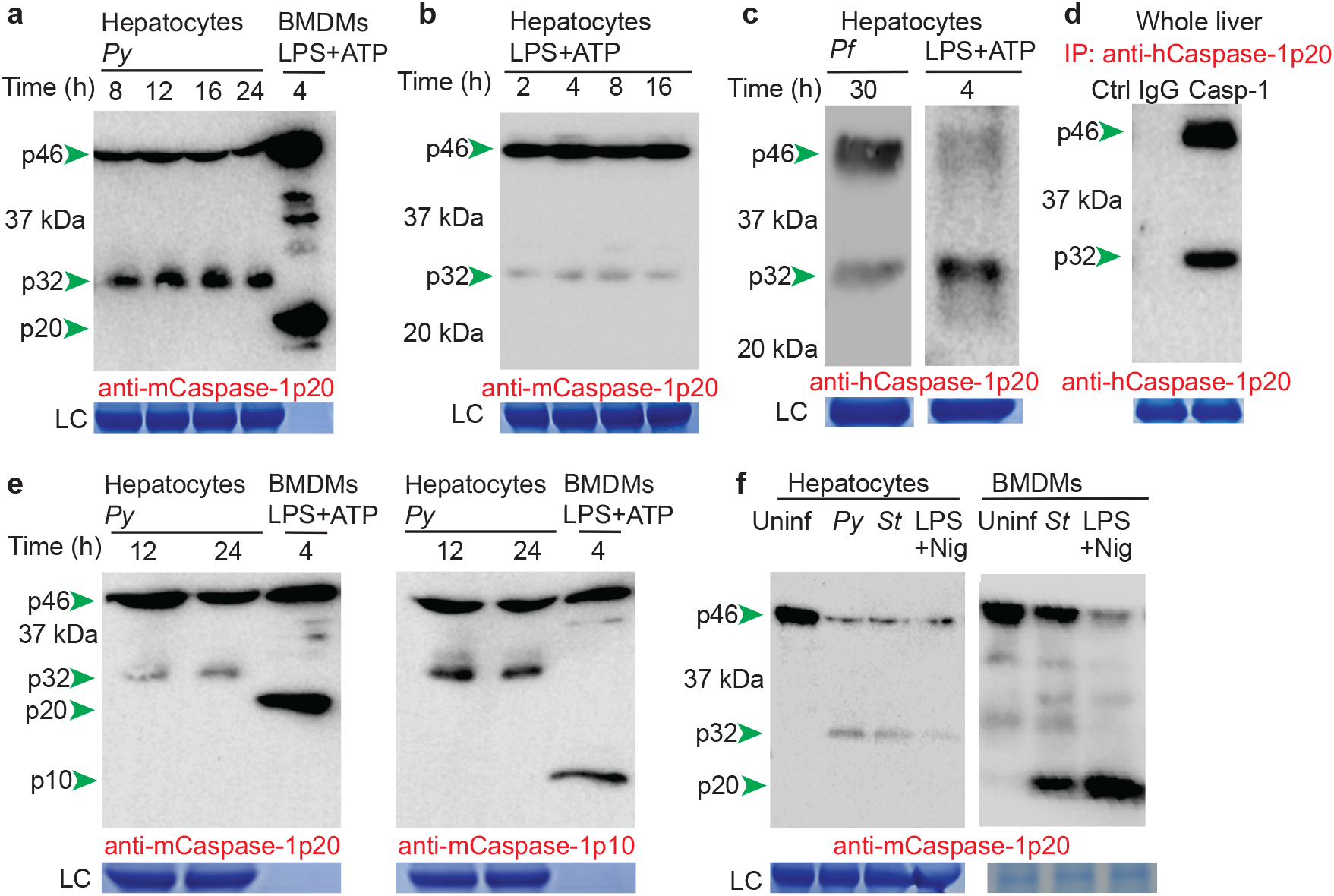
Incomplete cleavage and activation of Caspase-1 in hepatocytes. **a-b**, Immunoblot screen for Caspase-1 cleavage forms in primary mouse hepatocytes infected with *Py* (**a**) or treated with LPS+ATP (**b**) for the indicated time-frames. Murine BMDMs co-incubated with LPS+ATP served as the control. Caspase-1 p20 subunit specific antibodies detect uncleaved procaspase-1 (p46) and the cleaved Caspase-1 products, p32 or p20. **c**, Immunoblot analysis for cleaved Caspase-1 in human hepatocytes infected with *Pf* (left) or treated with LPS+ATP (right) for the indicated time-frames. **d**, Immunoblot analysis for Caspase-1 pulled down from whole-liver lysates of liver-humanized mice infected with *Pf* (30h p.i.), using with anti-p20 specific antibody. **e**, Immunoblot analysis for Caspase-1 cleavage in primary mouse-hepatocytes infected with *Py* and probed with Caspase-1 p20 subunit (left) or Caspase-1 p10 subunit (right) specific antibodies. Murine BMDMs co-incubated with LPS+ATP served as the standard for conventional Caspase-1 cleavage pattern, indicating p20 and p10. **f**, Immunoblot analysis for Caspase-1 cleavage in primary mouse hepatocytes (left) or BMDMs (right) co-incubated with *Py* (24h), *S. typhimurium* (*St*, 24h) or LPS + Nigericin (Nig, 4h). Uninfected cultures (Uninf) served as controls. In **a-f**, LC represents the protein loading controls from the SDS-PAGE, to provide an estimate of the representation of pro/Caspase-1 in the total protein content of the hepatocytes or BMDMs. Note that these are single exposure blots and due to the high signal intensity, the amount of total protein added from the BMDM lysates was insufficient for detection by Coomassie blue staining in the loading control. Data represent >3 independent experiments.

### ASC limits Caspase-1 cleavage in hepatocytes

A possible reason for the incomplete processing of Caspase-1 in hepatocytes is the presence of alternately spliced forms of procaspase-1, with their IDLs not amenable to proteolytic cleavage. However, RT-PCR followed by sequencing of uninfected, *Py* infected or LPS+ATP stimulated purified hepatocytes failed to indicate the presence of any alternate spliced procaspase-1 transcripts (Extended Data Fig. 11). In addition, bioinformatic analysis of our^10^ and others’^31–34^ published RNAseq data also showed no procaspase-1 variants in the infected or uninfected human or mouse hepatocytes. As discussed above, the recruitment of procaspase-1 to the ASC-speck is crucial to enable its inherent auto-proteolytic function through proximity-induced dimerization^7,17,35^. Merely maintaining recombinant procaspase-1 protein at high concentrations *in vitro* results in procaspase-1 achieving catalytically active quaternary structure and the ability to spontaneously self-cleave; both of which are lost upon adequate dilution^19^. Hence, we surmised that cleavage and activation of the constitutively expressed procaspase-1 in cells is facilitated by procaspase-1 achieving a critical local concentration in the cytosol, presumably at the inflammasome complex. Lower overall expression of procaspase-1 in hepatocytes could prevent it from achieving a sufficient density at the ASC speck, or the inadequate availability of ASC could preclude ASC oligomerization, leading to the generation of smaller specks and the recruitment of fewer procaspase-1 molecules^18,36^. Examination of publicly available transcriptional data^31–34^ and western-blot analyses indicated that both procaspase-1 and ASC are inherently less abundant in mouse and human hepatocytes, compared to in immune cells such as monocytes or macrophages (Fig. 4a, Extended Data Fig. 12a-b). This suggested that increasing expression of procaspase-1, ASC or both in hepatocytes could potentially induce complete autoproteolysis of procaspase-1. Indeed, transgenic over-expression of ASC resulted in the generation of Caspase-1 p20 following LPS+ATP stimulation or *Py* infection in primary hepatocytes (Fig 4b, Extended Data Fig 12c-e). Conversely, downmodulating ASC expression in BMDMs with ASC siRNA resulted in the formation of p32 in BMDMs (Fig 4c, Extended Data Fig 12 f-g). Taken together, these findings indicated that the expression level of ASC is a critical determinant of the extent of Caspase-1 cleavage in cells, and by extension, the nature of innate immune responses elicited by the cells.

**Figure 4:**
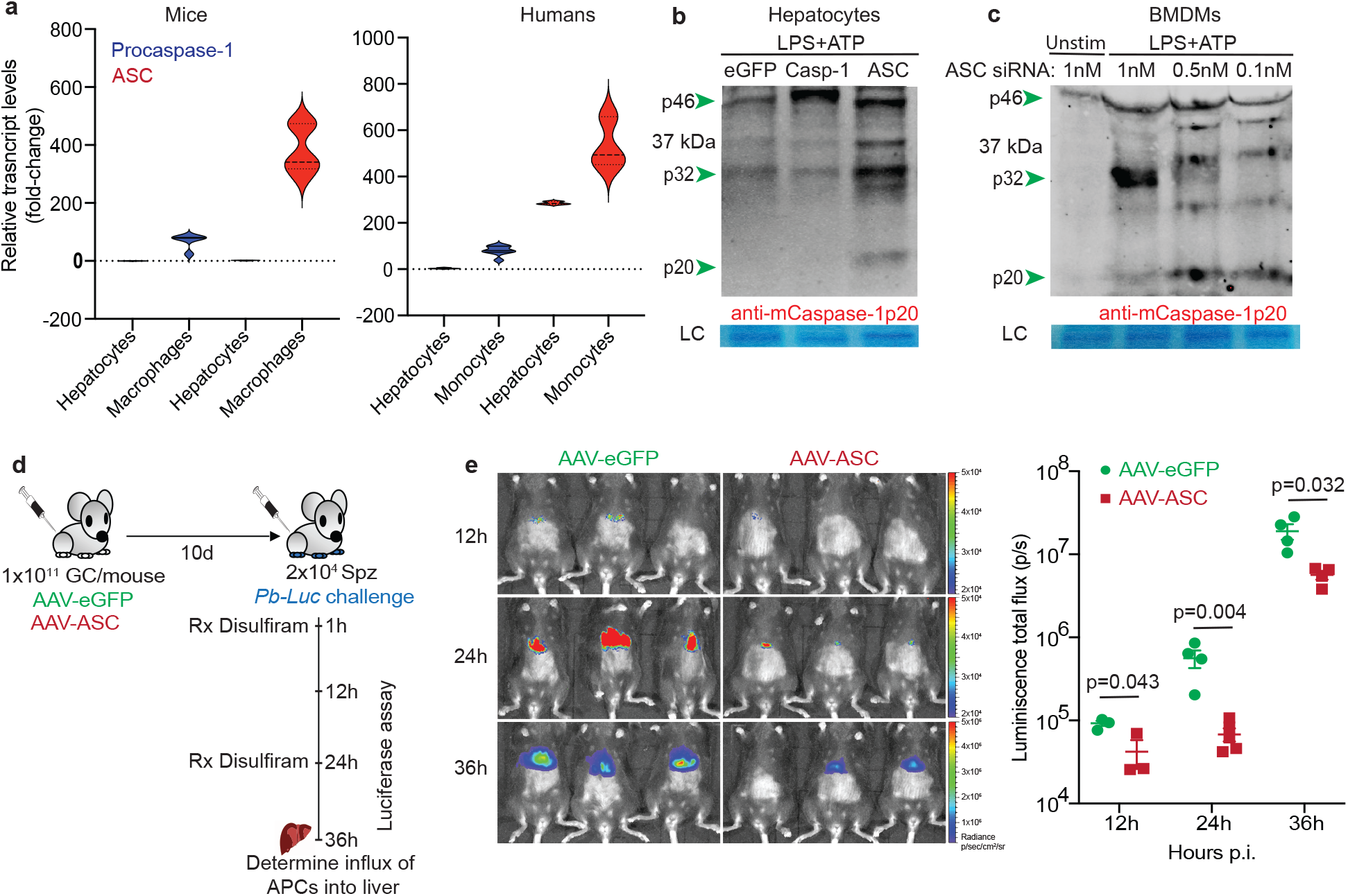
Inherently reduced expression of ASC in hepatocytes precipitate incomplete cleavage of Caspase-1, limited control of malaria infection. **a**, Combined data depicting the relative transcript levels of procaspase-1 and ASC in primary hepatocytes, macrophages or monocytes derived from mice or humans. **b**, Immunoblot analysis for Caspase-1 cleavage products in primary mouse hepatocytes transfected with mammalian expression vectors encoding eGFP, procaspase-1 (Casp-1) or ASC and stimulated in culture with LPS+ATP for 16h from 24h post transfection. **c**, Immunoblot analysis for Caspase-1 cleavage in BMDMs transfected with reducing dosages of ASC siRNA and stimulated with LPS+ATP for 4h at 24h post transfection. LC: Loading controls. **d**, Experimental scheme to determine the impact of ASC over-expression in hepatocytes, on the control of liver-stage malaria. 1×10^11^ genome copy of AAV-ASC or AAV-eGFP were inoculated into each B6 mice and challenged with 2×10^4^ sporozoites of *Pb-Luc* at d10 post inoculation. Mice were then imaged for total luminescence at the indicated time-points to determine the kinetics of parasite control in the liver. Livers collected at 36h p.i. to determine the infiltration of APCs into the liver **e**, Representative rainbow images of total luminescence in the livers of these mice at the indicated time points. Scatter plot in the right panel summarizes the data. Data presented as mean + s.e.m, analyzed with 2-tailed t-tests at each time-point to yield the presented p values. All data represent >2 independent experiments.

It is possible to generate p32 in BMDMs by mutating the autoproteolytic cleavage sites in the IDL sequence of procaspase-1^20^. Such BMDMs were unable to efficiently mature IL-1β in response to *St* infection or LPS+ATP stimulation^20^. Similarly, altering ASC through site directed mutagenesis to hamper the formation of ASC speck also impeded the maturation of IL-1β in BMDMs^18^. However, these alterations had a relatively low impact on GSDMD activation or the ability of BMDMs to undergo pyroptotic cell-death. Of note, procaspase-1 by itself can cleave GSDMD to induce pyroptotic cell-death, although fully cleaved Caspase-1 is more efficient in activating GSDMD^37^. This suggested that the extent of Caspase-1 cleavage may also govern GSDMD function. Hence, we hypothesized that complete processing of Caspase-1 facilitated by adequate ASC expression in hepatocytes would lead to efficient proteolytic maturation of IL-1 and IL-18, and improved GSDMD function. Hepatocytes over-expressing ASC were able to generate tangible levels of mature IL-1β and IL-18 (Extended Data Fig 13), supporting this hypothesis. The alternately cleaved Caspase-1 products observed in epithelial cells, neurons or some cancer cells^38–44^ might signal such nuances in Caspase-1 functions.

### Increasing ASC improves immunity to malaria

To ascertain the extent to which conventional activation of Caspase-1 in hepatocytes would impact immunity to malaria in the liver, we over-expressed ASC in the hepatocytes of mice. To enhance ASC expression specifically in the hepatocytes, we generated adeno associated viral vectors on AAV-DJ background encoding ASC (AAV-ASC) or control eGFP (AAV-eGFP), under the hepatocyte-specific albumin promoter^45–48^. Mice inoculated with AAV-ASC or AAV-eGFP were challenged with *Pb* expressing luciferase (*Pb-Luc*) to determine the kinetics of *Plasmodium* infection in the liver (Fig. 4d). Transgenically enhancing ASC expression in hepatocytes also aided in a more rapid control of *Plasmodium* infection in mice (Fig 4d-e, extended Data Fig 14a). Over-expression of ASC in hepatocytes enhanced their ability to undergo cell-death in response to *Py* infection or LPS+ATP stimulation (Extended Data Fig. 14b). Disulfiram treatment negated the enhanced cell-death observed in hepatocyte in response to LPS+ATP treatment (Extended Data Fig 14c), indicating that the enhanced hepatocyte cell death promoted by ASC overexpression was indeed mediated by GSDMD. Treatment with disulfiram also prevented the improved control of *Plasmodium* infection observed in mice over-expressing ASC in hepatocytes (Extended Data Fig. 14d). Transgenically enhancing ASC expression in hepatocytes also resulted in increased influx of CD11c^+^ and CSF1R^+^ CD11c^+^ antigen presenting cells (APCs) into the liver following *Plasmodium* infection (Extended Data Fig. 15), possibly driven by IL-1 or IL-18^49,50^. Although not pertinent to the direct control of an ongoing infection, these infiltrating APCs are vital for the generation of subsequent adaptive immune responses and vaccine-induced immunity to malaria^10^. Taken together, our data indicated that enhancing ASC expression induced conventional cleavage of Caspase-1 in hepatocytes in response to *Plasmodium* infection, possibly leading to more efficient GSDMD mediated elimination of the infected hepatocytes, precipitating better control of malaria infection in the liver.

Our discoveries offer novel insights into the mechanism by which *Plasmodium* parasites interact with and elicit host innate defenses in hepatocytes. The incomplete processing of Caspase-1 in hepatocytes, which severely limits inflammatory cytokine production and appears to elicit suboptimal pyroptotic cell-death, may have contributed to the liver being uniquely exploitable for *Plasmodium* infection and survival. As a frontline defensive organ that drains the gastrointestinal tract replete with potential pathogens and toxins, the ability of the liver to mount adequate immune responses without overreacting is crucial for the survival of the host^51^. The incomplete processing of Caspase-1 in hepatocytes may be an evolutionary adaptation to restrict the pro-inflammatory consequences of the inflammasome pathway without renouncing the ability to combat pathogens like *Plasmodium* through programmed cell-death.

## Acknowledgements

We thank Rick Tarleton and Ronald Etheridge for comments, Gibran Nasir for preparing humanized mice for analyses, Magdy Alabady for assistance with scRNAseq, Teneema Kuriakose for helpful discussions and Fayyaz Sutterwala for mice, the UGA CTEGD Flow Cytometry Core, UI Central Microscopy Research facility, UGA CTEGD Sporocore, UGA Georgia Genomics and Bioinformatics Core, Iowa Institute of Human Genetics, UGA and UIowa animal research facility staff and the NYU and Johns Hopkins Malaria Institute Insectary Cores. Support for these studies was provided by NIH (AI85515, AI95178, AI100527 to JTH, AI132359 to PS and K22AI127836 to PG) and the UGA Research Foundation (Startup funding to SPK).

## Author contributions

CMdS, BP, RPB, KP, JCS, PS, PG, JTH and SPK designed the experiments. CMdS, BP, RPB, KP, LSH, LLP, MJS and SPK performed the experiments. SPK and JTH wrote the manuscript. DEK and T-DK provided vital resources for the study.

## Competing interests

The authors declare no competing interests.

## Notes

### Competing Interest Statement

The authors have declared no competing interest.

