## Supplementary Data for "Altered cleavage of Caspase-1 in hepatocytes limits control of malaria in the liver"

### Supplementary Figures

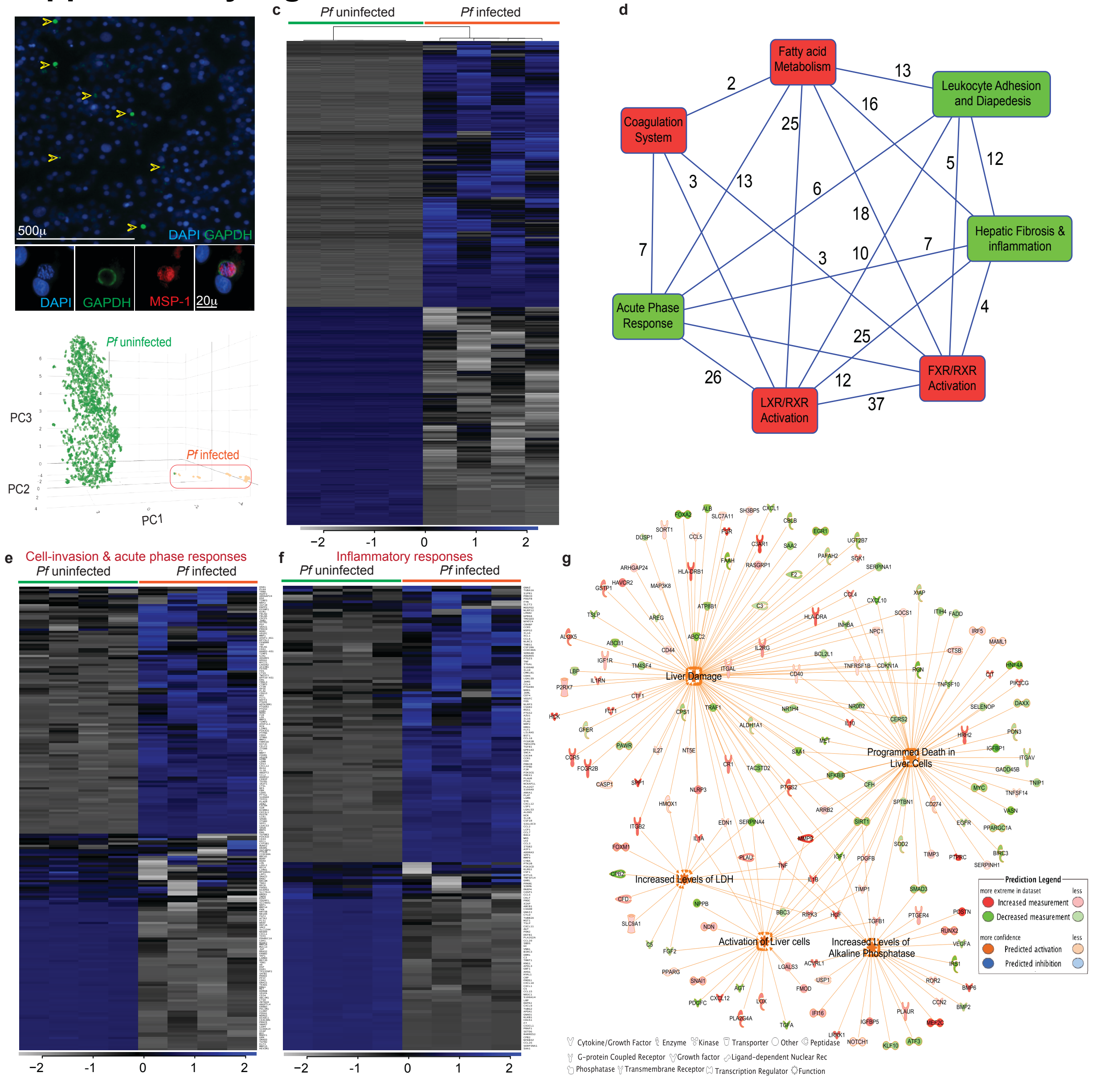


**Extended data Figure 1: Transcriptional upregulation of inflammatory cell-death pathways in *P. falciparum* infected hepatocytes. a,** Representative pseudo-colored fluorescent micrographs showing in vitro cultured primary human hepatocytes infected with *Pf,* 4d p.i. The individual panels show staining with separate *Pf*-specific antibodies (GAPDH and MSP-1) or DNA (DAPI), and fluorescence overlays. **b-c**, Transcriptional differences determined by single-cell RNA sequencing of *Pf* infected or uninfected primary human hepatocytes. *Pf* infected hepatocytes were identified by the presence of *Pf*-specific transcripts, which corroborated the unsupervised clustering of hepatocytes based on gross transcriptional changes. Principal component analysis (**b**) and heat map (**c**) representing gross transcriptional changes observed between the infected and uninfected hepatocytes. **d**, Top canonical biochemical pathways perturbed by *Pf* infection in primary human hepatocytes (4d p.i.), determined by gene-set enrichment analysis through the gene ontology algorithms of Ingenuity Pathway Analysis (IPA). The connecting lines and the corresponding numbers indicate overlapping genes. The canonical pathways pertinent to host immune responses are indicated in green. **e-f**, Transcriptional differences in the indicated genes that are components of cell-invasion and acute phase response (**e**) and inflammatory response (**f**) pathways, determined in human hepatocytes infected with *Pf*, 4d p.i. **g**, Significantly enriched network of functional interactions evaluated by IPA comparing *Pf* infected and uninfected human hepatocytes. The nodes represent the key factional outcomes predicted based on the transcriptional identity (represented by color coded proteins, see legend) of the *Pf* infected primary human hepatocytes at 4d p.i.

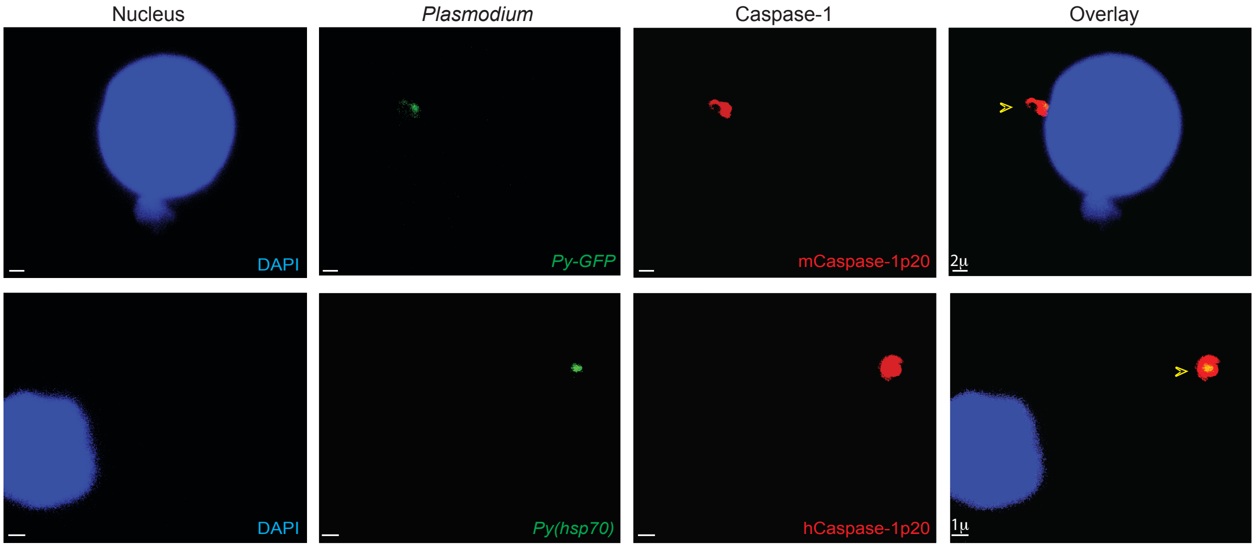


**Extended Data Figure 2: Localization of Caspase-1 in *Plasmodium* infected hepatocytes.** Representative pseudo-colored confocal image from frozen liver-sections showing Caspase-1 in *Py* infected murine (upper panel, 16-20hpi) or *Pf* infected human (lower panel, from liver-humanized mice, 30hpi) hepatocytes. Data presented represents >3 separate experiments.

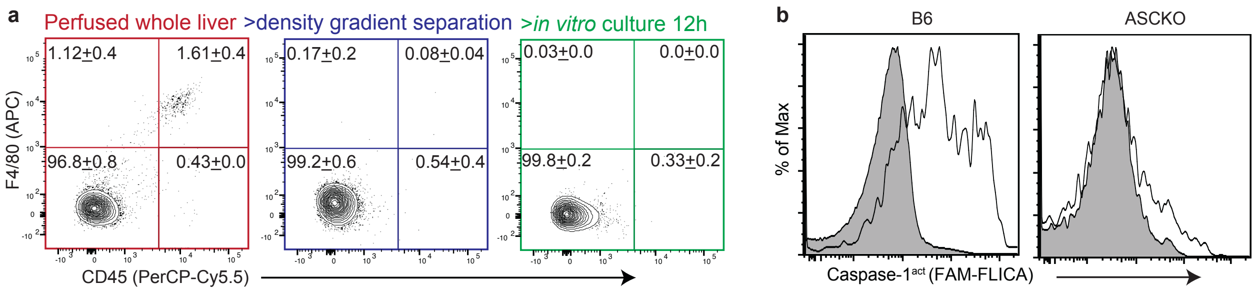


**Extended Data Figure 3: Caspase-1 activation in hepatocytes depend on ASC. a**, Flow-plots examining the presence of contaminating hematopoietic ($CD45^+$) or Kupffer ($F4/80^+$ $CD45^+$) cells in the perfused whole liver single-cell suspension (left panel), post density gradient purification of hepatocytes (middle panel) and after subsequent ex vivo culture for 12h (right panel), during the process of generating primary hepatocytes from mice. Numbers inset in quadrants represent frequencies as mean ± s.e.m. Data represents more than 5 separate experiments. **b**, Histograms depicting Caspase-1 activation $tdTomato^+$ *P. yoelii* infected B6 or AscKO primary hepatocytes at 30hpi. Grey histograms represent the uninfected ($tdTomato^-$) hepatocytes in the same culture.

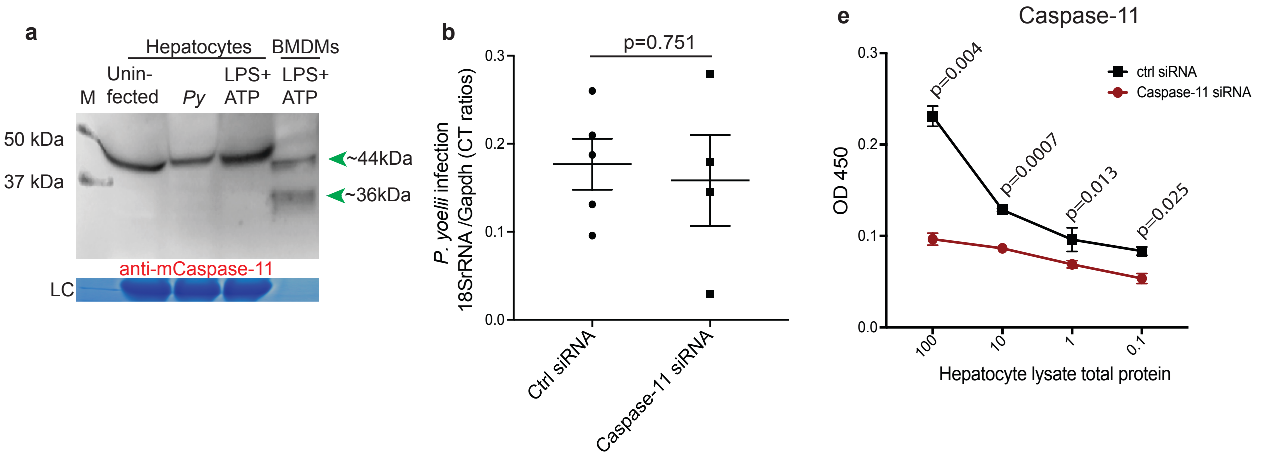


**Extended Figure 4: Caspase-11 not induced by *Plasmodium* infection in hepatocytes and do not aid in the control of liver-stage malaria. a,** Immunoblot analysis for Caspase-11 cleavage in primary B6 hepatocytes co-incubated with *Py* (24h) or LPS+ATP. LC: loading control. **b**, Scatter plots showing relative liver-parasite burdens, 36h p.i. in B6 mice that received control or Caspase-11 siRNAs by hydrodynamic injection 48h prior to *Py* inoculation i.v. Dots represent individual mice, data presented as mean + s.e.m and analyzed by 2-tailed t-test yielding the indicated p value. **c**, Relative protein levels determined by ELISA in the primary hepatocyte lysates derived from B6 mice at 48h after inoculation with the indicated siRNAs by hydrodynamic injection and LPS+PolyI:C at 24h. Data represent 1 of 2 separate experiments with 3 mice/ group, compared with 2-tailed t-tests at each dilution across biological replicates to yield the indicated p values. All data presented represents at least 3 separate experiments.

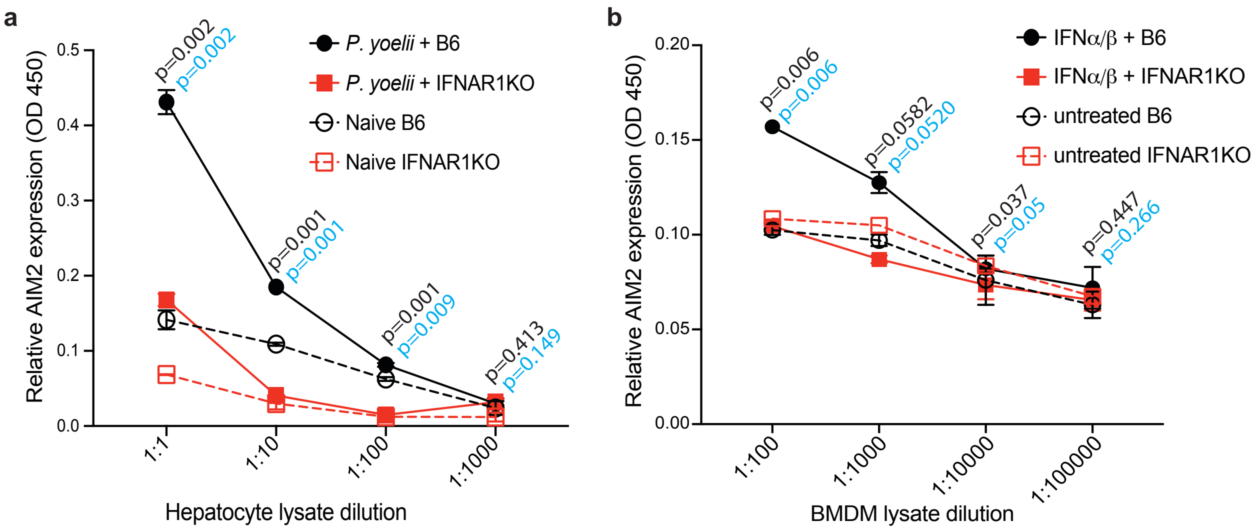


**Extended Data Figure 5: *Plasmodium* infection induces type-1 IFN dependent AIM2 expression in hepatocytes. a**, Relative AIM2 expression determined by ELISA in hepatocytes derived from both B6 or Ifnar1KO mice, infected with *Py,* 24h p.i. Data presented as mean $\pm$ s.e.m analyzed with t-tests at each dilution of total hepatocyte lysates. *Py* infected B6 and Ifnar1KO hepatocytes are compared, yielding the p values indicated in black or *P. yoelii* infected or uninfected B6 hepatocytes are compared yielding the p values indicated in blue. **b**, Relative AIM2 expression determined by ELISA in BMDMs derived from both B6 or Ifnar1KO mice, treated with IFN$\alpha$+IFN$\beta$ for 24h. Data presented as mean $\pm$ s.e.m analyzed with t-tests at each dilution of total BMDM lysate, comparing IFN$\alpha$+IFN$\beta$ treated B6 and Ifnar1KO BMDMs yielding the p values indicated in black or IFN$\beta$ treated or untreated B6 BMDMs yielding the p values indicated in blue. Data represents 1 of 3 separate experiments.

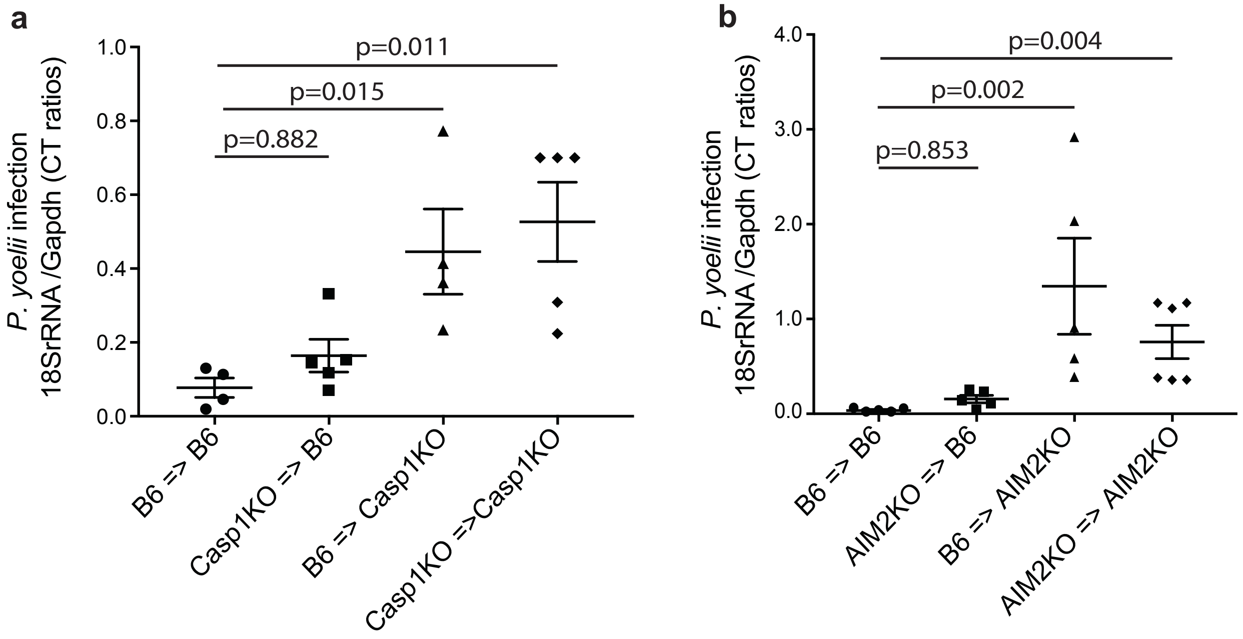


**Extended Data Figure 6: Myeloid cells do not aid in Caspase-1 mediated control of malaria in the liver. a**, Scatter plots showing relative parasite burdens in the whole liver at 36h p.i. in B6 or Casp1KO chimeric recipient mice reconstituted with B6 or Casp1KO bone-marrow and inoculated with *Py* i.v. **b**, Scatter plots showing relative whole liver-parasite burdens at 36h p.i. in B6 or AIM2KO chimeric recipient mice reconstituted with B6 or AIM2KO bone-marrow and inoculated with *Py* i.v. The dots in the scatter plots represent individual mice, data presented as mean $\pm$ s.e.m and analyzed by ANOVA with Dunnett's corrections, yielding the indicated p values, and is representative of $\geq$3 separate experiments.

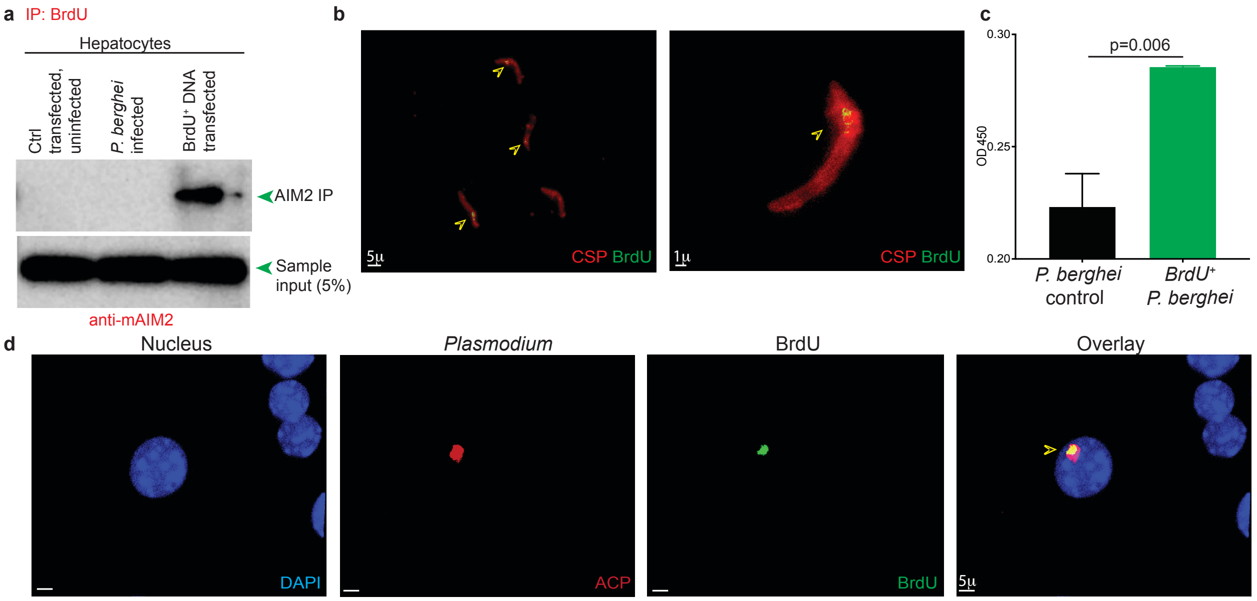


**Extended Data Figure 7: DNA associates with AIM2 in hepatocytes, BrdU incorporated into *Plasmodium* DNA. a**, Immunoblot analysis for AIM2 co-immunoprecipitated with anti-BrdU antibodies, from whole-cell lysates of mock transfected, $BrdU^{-}$*Pb* infected or control BrdU-incorporated B16 melanoma cell DNA transfected cultured primary hepatocytes, 24h post infection/ transfection, as a proof of concept for Fig 1f. **b**, Pseudo-colored confocal image of a representative field of $BrdU^{+}$ *Pb* (anti-Circumsporozoite protein, CSP) sporozoite stages derived from infected mosquitoes. Arrows indicate BrdU incorporation. **c**, ELISA comparing lysates of *Pb* sporozoites derived from BrdU-fed or control infected mosquitoes to determine relative BrdU levels. Data presented as mean $\pm$ s.e.m from triplicate readings, compared with t-tests to yield the presented p values. **d**, Representative pseudo-colored confocal images of a $BrdU^{+}$ *Pb* (anti-Acyl carrier protein, ACP) merozoite stage in the liver of B6 mouse at 24h p.i. Arrow indicates the developing parasite. All data represent at least 3 separate experiments.

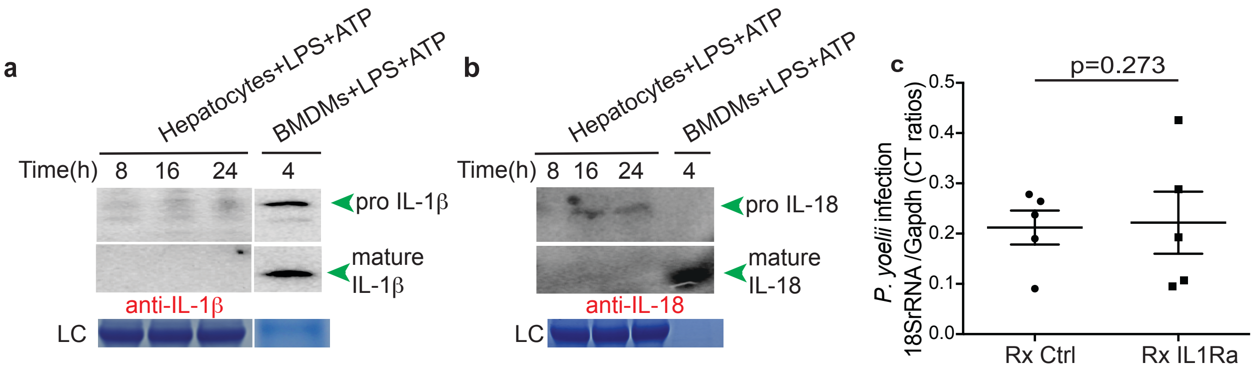


**Extended Data Figure 8: Caspase-1 activation does not induce mature IL-1β, IL-18 in hepatocytes. a-b**, Immunoblot analysis for IL-1β (**a**) and IL-18 (**b**) in supernatants of primary B6 mouse hepatocytes co-cultured with *Py* for 24h. LPS+ATP co-incubated with primary hepatocytes or BMDMs from B6 mice for the indicated lengths of time served as controls. LC: loading control. **c**, Scatter plots indicating relative liver-parasite burdens at 36h p.i. in B6 mice treated with IL-1R antagonist, Anakinra or the vehicle control (-1,0,1dpi) and inoculated with *Py* i.v. Dots represent individual mice, data presented as mean $\pm$ s.e.m and analyzed with 2-tailed t-test yielding the indicated p value. All data represent at least 2 separate experiments.

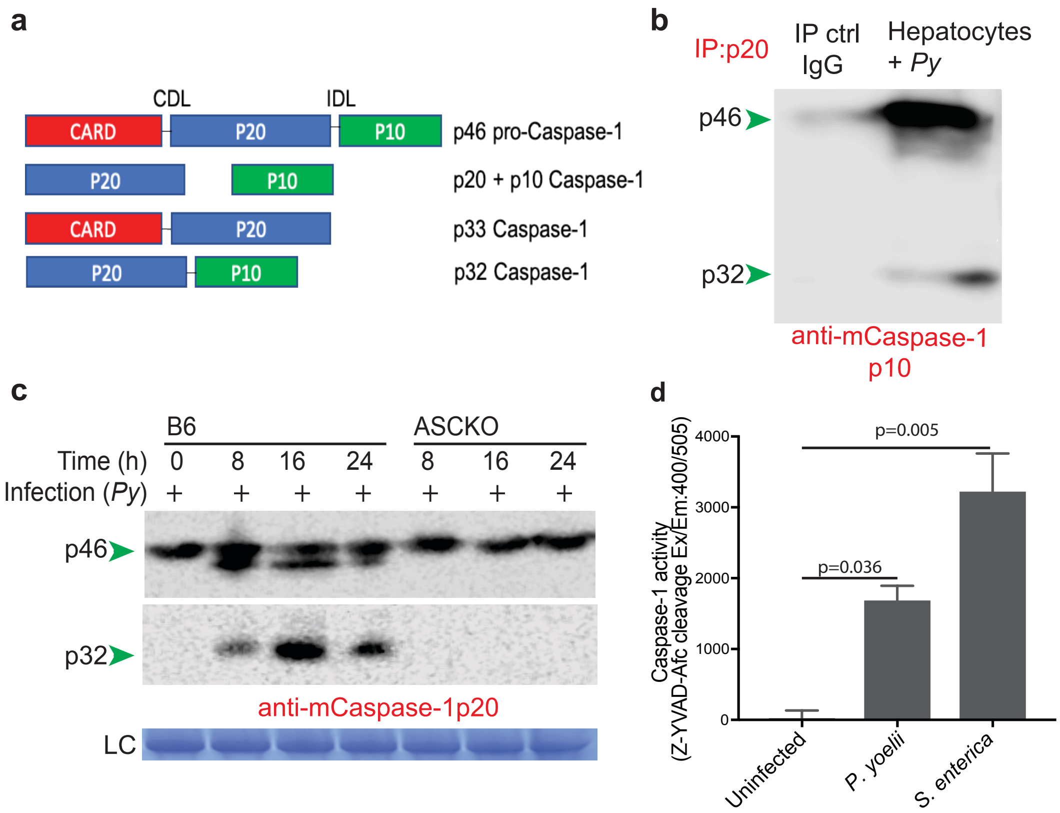


**Extended Data Figure 9: p32 in hepatocytes composed of p20 and p10, ASC dependent, and catalytically active. a,** Current understanding of Caspase-1 activation: Autoproteolytic processing of 46kDa procaspase-1 (p46) into a catalytically active heterotetramer composed of p20 and p10 domains. A very short-lived p33 intermediate composed of CARD and p20 has been reported in macrophages. We observe p32 composed of unseparated p20 and p10 in hepatocytes. CDL: CARD domain linker, IDL: Interdomain linker. **b**, Immunoblot analysis in whole-cell lysates of primary B6 hepatocytes co-cultured with *Py* for 24h, immuno-precipitated with anti-mCaspase-1p20 and the precipitate probed with anti-mCaspase-1p10. **c**, Immunoblot analysis for Caspase-1 cleavage in B6 or ASCKO mice hepatocytes co-incubated with *Py* for the at the indicated times. LC: Loading control. **d**, Caspase-1 proteolytic activity measured by the cleavage of Z-YVAD-Afc. Caspase-1 obtained from *Py* infected B6 primary hepatocyte cultures at 24h p.i. Uninfected or *S. enterica* infected hepatocytes served as controls. Data from represents 3 biological replicates/group, presented as mean $\pm$ s.e.m, analyzed by ANOVA with Tukey's correction to provide the indicated p values. All Data presented represents at least 2 separate experiments.

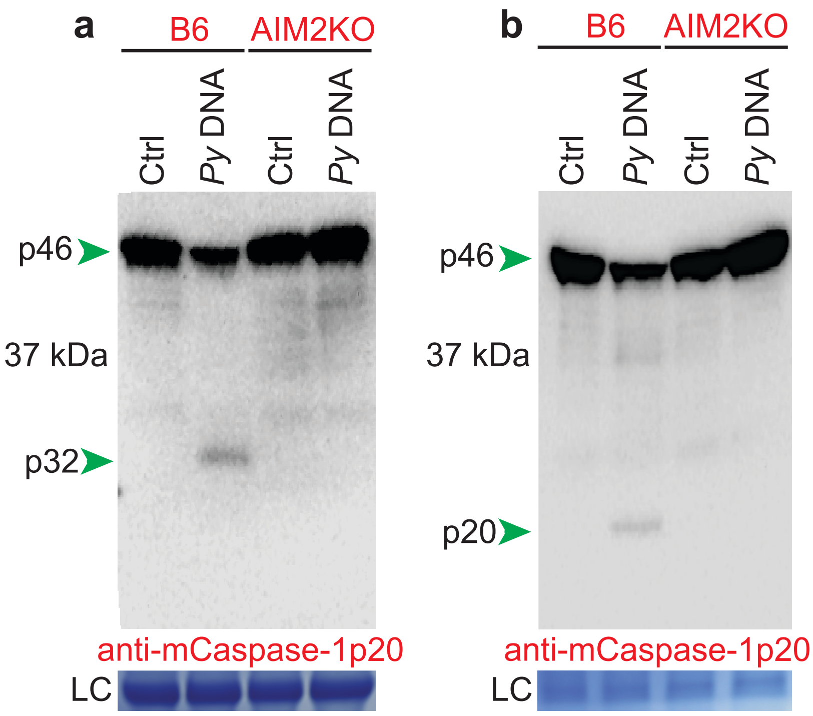


**Extended Data Figure 10: *Plasmodium* DNA induces AIM2 mediated incomplete cleavage of Caspase-1 in hepatocytes.** Immunoblot analysis for Caspase-1 cleavage in primary hepatocytes (**a**) or BMDMs (**b**) derived from B6 or AIM2KO mice and transfected with *Py* DNA after pretreatment with LPS for 4h. Whole cell lysates collected 24h post transfection. LC: loading control. Data represents one of 3 separate experiments.

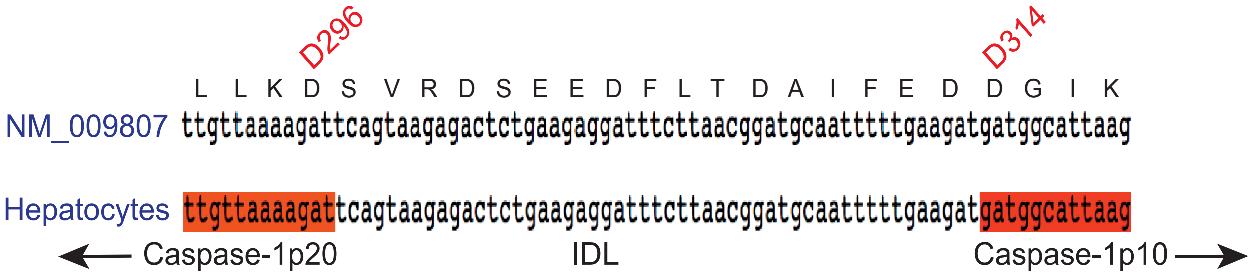


**Extended Data Figure 11: Sequence of the IDL region of procaspase-1 in B6 mice hepatocytes.** Sequence of p20-p10 IDL and flanking regions of procaspase-1 transcripts from B6 hepatocytes compared to the reference sequence. Represents the sequence from >15 separate sequencing clones. D296 and D314 indicates the known autoproteolytic cleavage sites in the Caspase-1 isoform of murine BMDMs.

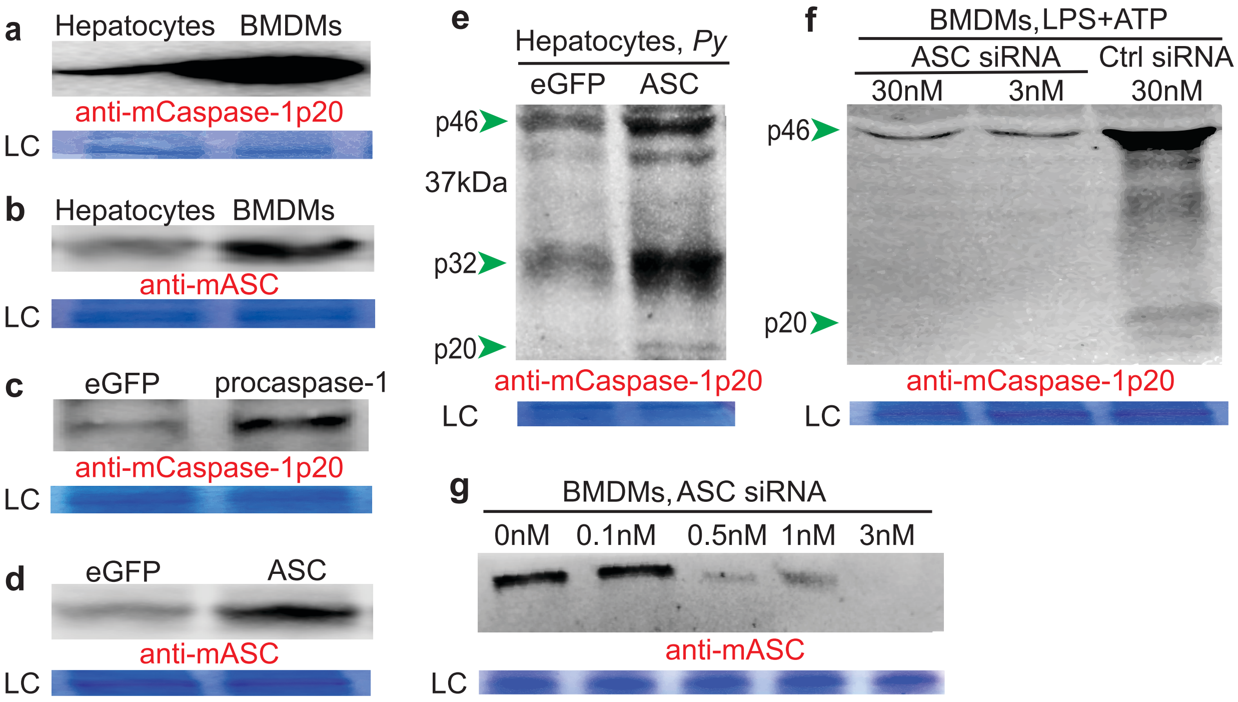


**Extended Data Figure 12: ASC expression in cells impact Caspase-1 processing. a-b,** Immunoblot analysis for the relative expressions of procaspase-1 (**a**) or ASC (**b**) in resting primary mouse hepatocytes or BMDMs**. c-d,** Immunoblot analysis for the relative expressions of procaspase-1 in hepatocytes transfected with mammalian expression plasmids encoding eGFP or procaspase-1 (**c**) or of ASC in hepatocytes transfected with mammalian expression plasmids encoding eGFP or ASC (**d**). **e**, Immunoblot analysis for Caspase-1 cleavage in primary mouse hepatocytes transfected *in vitro* with control eGFP or ASC encoded mammalian expression plasmids and subsequently (16h post transfection) infected with *Py*, 24h p.i. **f**, Immunoblot analysis for Caspase-1 cleavage in BMDMs transfected with different doses of ASC siRNA and stimulated with LPS+ATP (4h) indicating no Caspase-1 processing following complete loss of ASC expression. **g**, Immunoblot analysis for the expression of ASC in BMDMs transfected with the indicated dosages of ASC siRNA to downregulate ASC expression. LC: loading control. All data presented represent one of at least 3 separate experiments.

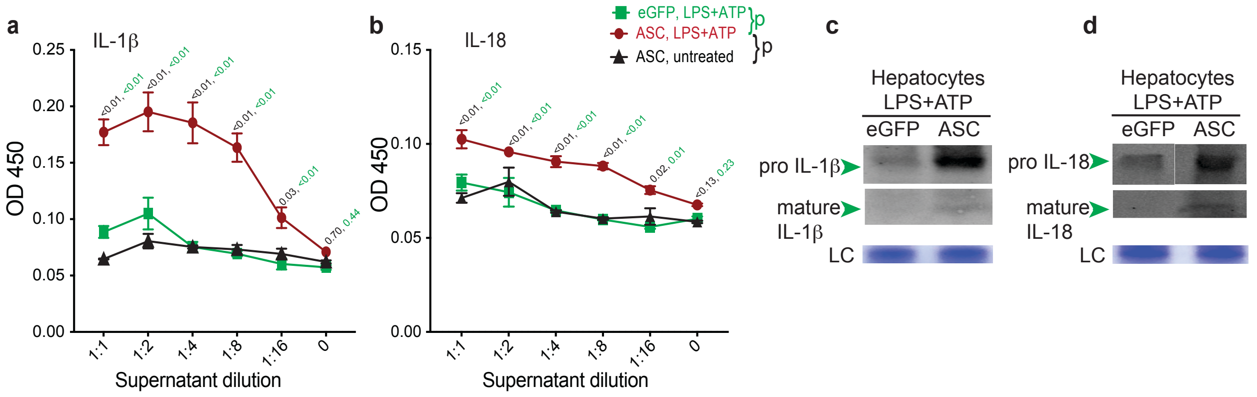


**Extended Data Figure 13: Enhancing ASC expression facilitates IL-1b and IL-18 production in hepatocytes. a-b**, Relative levels of IL-1β (**a**) or IL-18 (**b**) in the supernatants of primary mouse hepatocytes transfected with control eGFP or ASC encoding mammalian expression plasmids and treated 24h later with LPS+ATP for 24h. Data presented as mean ± s.e.m at each dilution of the culture supernatant, analyzed using 2-way ANOVA with Dunnett's correction to yield the p-values comparing the groups as indicated by color codes. **c-d**, Immunoblot analysis for IL-1β (**c**) or IL-18 (**d**) in supernatants of primary B6 mouse hepatocytes treated with LPS+ATP for 24h. LC: loading control. All data represents one of 3 separate experiments.

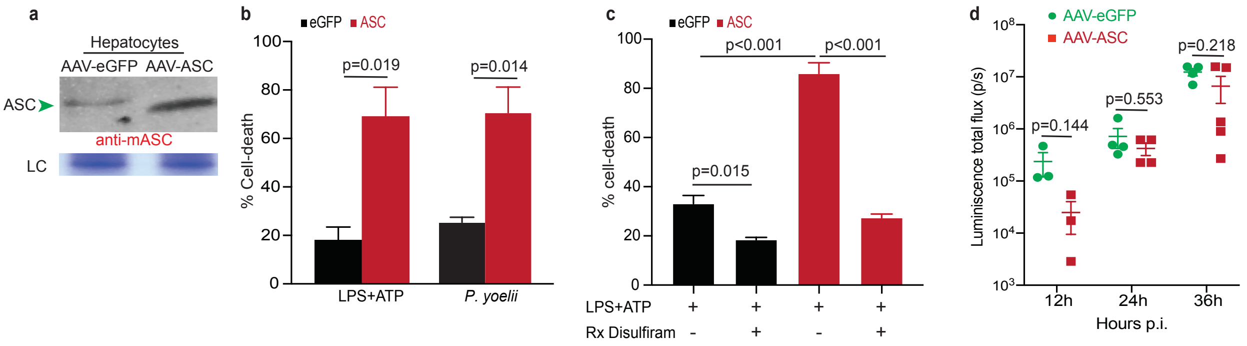


**Extended Data Figure 14: Enhancing ASC expression leads to increased GSDMD mediated pyroptotic cell-death in hepatocytes. a,** Immunoblot analysis for the relative expressions of ASC in hepatocytes derived from mice inoculated with AAV-eGFP or AAV-ASC at 10d p.i. LC: loading control. **b**, Relative cell-death determined by PI staining in hepatocytes transfected with mammalian expression plasmids encoding eGFP or ASC and treated with LPS+ATP (16h) or co-incubated with *Py* (16h). **c**, Relative cell-death determined by PI staining in hepatocytes transfected with mammalian expression plasmids encoding eGFP or ASC and treated with LPS+ATP or disulfiram (16h). **d**, Scatter plots showing relative liver-parasite burdens at the indicated time points in mice inoculated with AAV-eGFP or AAV-ASC and at 10d p.i challenged with *Pb-Luc* while under disulfiram treatment (please see Fig 4d). Data presented as mean $\pm$ s.e.m and analyzed using 2-tailed t-tests (b, d) or ANOVA with Tukey's correction (c), yielding the indicated p values. All data representative of $\geq$2 separate experiments.

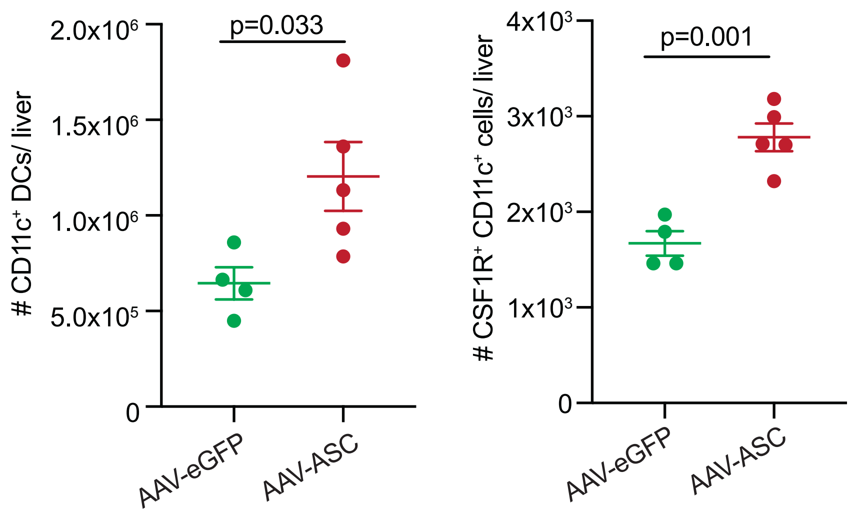


**Extended Data Figure 15: ASC over-expression augments CD11c$^+$ and CSF1R$^+$ APC infiltration to the liver following *Plasmodium* infection.** Scatter plots depicting the total numbers of CD11c$^+$ (left) and CSF1R$^+$ CD11c$^+$ (right) APCs in the liver following *Pb-Luc* challenge (36h p.i.) of mice inoculated with AAV-eGFP or AAV-ASC as depicted in Fig 4d. Data presented as mean $\pm$ s.e.m and analyzed using 2-tailed t-tests, to yield the indicated p values. Represents 2 separate experiments.

**Supplementary Video S1: *Plasmodium* infected hepatocyte undergoes pyroptotic death**. Confocal time-lapse live-microscopy images showing *P. yoelii* (CellTrace Violet$^{+}$) infected primary hepatocytes in culture (tdTomato$^{+}$), imaged from 24-32h of co-culture, at 0.5h intervals. Infected cells indicated by arrowheads.

**Supplementary Video S2: pyroptotic death in *Plasmodium* infected hepatocytes is dependent on GSDMD**. Confocal time-lapse live-microscopy images showing *P. yoelii* (GFP$^{+}$, green) infected primary hepatocyte in culture (CellTrace Violet$^{+}$, blue), imaged from 8-47h of co-culture, at 0.5h intervals. Infected cell indicated by the arrowhead.

#### Supplementary methods

##### Mice and pathogens

C57BL/6 (B6) mice were purchased from the National Cancer Institute or Jackson Laboratory and liver-humanized mice were obtained from Yecuris. GsdmdKO mice were provided by Dr. Thirumala-Devi Kanneganti, tdTomato expressing mice by Dr. John Englehardt (University of Iowa), ASCKO, NLRP3KO, NLRC4KO and Casp1KO mice by Dr. Fayyaz Sutterwala (Cedar-Sinai), IFNAR1KO, NLRP1bKO and AIM2KO mice procured from the Jackson laboratory. All mice were housed with appropriate biosafety containment at the animal care units at the University of Iowa, Johns Hopkins University or the University of Georgia. The animals were treated and handled in accordance with guidelines established by the respective Institutional Animal Care and Use Committees. *Anopheles stephensi* mosquitos parasitized with *P. yoelii* 17XNL (*Py*) and GFP$^+$ *P. yoelii* 17XNL (*Py-GFP*) were obtained from New York University. *A. stephensi* mosquitos infected with *P. falciparum* NF54 or BD007 isolates were maintained at the insectaries at Johns Hopkins University or the University of Georgia. *A. stephensi* mosquitos infected with *P. berghei ANKA* (*Pb*) and *Pb* expressing luciferase (*Pb-Luc*) were reared at the University of Georgia insectary. *Salmonella enterica* serovar Typhimurium was cultured in tryptic soy broth (ThermoFisher) at 37°C to infect cells at an MOI of 5.

##### Primary mouse hepatocyte culture, *in-vitro* sporozoite infection and PRR stimulation.

Primary hepatocytes were isolated from mice as described in detail before[1]. In short, in anesthetized (Ketamine/Xylazine (87.5/12.5 mg/Kg)) mice, the inferior vena cava was catheterized (BD auto guard, 22G) aseptically to perfuse the liver by draining through the portal vein. A steady-state perfusion of the liver was performed first with PBS (4ml/ min for 5 minutes), then Liver Perfusion Medium (4ml/min for 3 minutes, Gibco), and finally Liver Digest Medium (4ml/ minute for 5 minutes, Gibco). Digested liver was excised, single cell suspension made and resuspended in a wash solution of 10% FCS (Sigma-Aldrich) in DMEM (Gibco). Hepatocyte fraction was recovered by centrifugation at 57g, from which debris and dead cells were then removed by density gradient centrifugation with a 46% Percoll (GE Healthcare) gradient. Remaining cells were counted and resuspended in DMEM with 10% FCS. $6x10^4$ cells were plated on flat bottomed collagen-coated plates and incubated at 37°C, 5% $CO_2$. $8x10^5$ primary mouse or human (obtained from BioIVT) hepatocyte cultures were infected with $2\text{-}4x10^4$ sporozoites in each well of a 6-well plate for reference. Cultures were further incubated for the desired time to allow infection and liver-stage parasite development. Hepatocytes were stimulated with LPS (100 ng for 3.5-15.5h as indicated, Invivogen), followed by ATP (5mM for 0.5h, Sigma) or nigericin (1mM for 0.5h, Alfa Aesar). $3x10^6$ BMDMs were treated with LPS (3.5h) followed by ATP (0.5h) to serve as controls, as described in detail before[2]. Culture supernatants and/or cell lysates were obtained at various time points post infection or stimulation as indicated.

**Primary human hepatocyte culture and *Pf* infection**

Primary human hepatocyte infection in this study followed published methods[3]. Briefly, cryopreserved primary human hepatocytes (BioIVT, NY, USA) were thawed two-days prior to infection and seeded with $18x10^3$ cells per well in a 384-well collagen-coated plate (Greiner). Hepatocytes was maintained with daily media change using customized InVitroGro HI medium devoid of dexamethasone (BioIVT), but supplemented with 5% human serum (Interstate Blood Bank), penicillin, streptomycin, neomycin and gentamicin (Gibco). The salivary glands of *Pf* BD007 (Bangladesh isolate) infected *A. stephensi* mosquitoes were collected, total sporozoite counts determined and $6x10^3$ sporozoites were inoculated into the hepatocyte culture. The Hepatocytes in culture were maintained at 38°C with 5% $CO_2$.

At 4d p.i., infection rates were determined by immunofluorescence assay. Infected hepatocytes were fixed with 4% paraformaldehyde (Alfa Aesar) for 10 min at room temperature and washed twice with PBS followed by permeabilization and blocking with 0.03% Triton-X (Acros) and 1% BSA (w/v) (Fisher Scientific) in PBS. The samples were concurrently stained with anti-GAPDH and anti-MSP antibodies (European Malaria Reagent Repository, UK) at 4°C overnight followed by staining with fluorescently labelled secondary antibodies at 4°C, overnight. The wells were then stained with DAPI under room temperature for 1 hr. The *Pf* infection in each well were imaged using ImageXpress and quantified using the MetaXpress software (Molecular Devices).

**Single-cell RNAseq and analysis**

Primary human hepatocytes infected with *Pf* were washed with PBS and the adherent hepatocytes were removed after trypsinization at 4d p.i. The detached cells were washed, counted, centrifuged at 57g for 3 minutes, supernatant removed and resuspended to a final concentration of 700 cells/μl in PBS. Subsequently scRNA-seq analysis was performed at the Georgia Genomics and Bioinformatics core at the University of Georgia in four replicate samples. Illumina NexSeq X10 RNAseq single cell paired end reads were generated and each single-cell fastq data was then submitted to CellRanger 3.0.1 (10Xgenomics). CellRanger counts were run using the human GRCh38-3.0.0 transcriptome as reference for each replicate. The resultant data were then submitted to a quality control filter to select cells with >1000 UMI counts/reads, >1000 detected genes, <20% of mitochondrial content and reads with >80% protein coding fraction. All ribosomal genes were also filtered out to streamline the transcriptional analysis. To remove noise in the acquired data, highly variable genes with False Discovery Rate of >0.1 and minimum dispersion of 0.5 were selected to identify the *Pf* infected or non-infected clusters. Mean-variance relationships of the log-counts were used for normalization by Voom[4] implemented in R. Limma Bioconductor package[5] for covariate removal of potential mitochondrial or ribosomal transcripts from the filtered dataset using linear regression. The final filtered data was then scaled using Seurat 3.0[6]. For three-dimensional reduced visualization of the cells, uniform manifold approximation and projection (UMAP) algorithm was used. The observed clusters were then predicted using the K-means with the Hartigan-Wong algorithm and 2:20 number of clusters to test. After getting the clusters we selected the barcodes represented in each one of them, and divided them in different files for the differential expression analysis. Subsequently, the DEseq2 Bioconductor package[7] was used to calculate the differential expression between the clusters. The *Pf* infected and uninfected clusters of hepatocytes were identified using the *Pf* transcript associated barcodes in the single cell data and their distribution among the clusters. This correlated with the expected rate of infection based on the experimental data generated by microscopic examination of concurrent *Pf* infected cultures. Visualization of the differentially expressed genes represented as heat maps were generated using heatmap3 R software package and the gene set enrichment and interaction pathways were generated using the Ingenuity Pathway Analysis software (Qiagen Bioinformatics).

**Determining differential transcription**

Distinct datasets were built for mice and humans. Mice RNAseq Illumina reads were retrieved from NCBI Sequence read archive (SRA) under accessions SRR9961636, SRR9961637, SRR9961638, SRR9961644, SRR9961945, and SRR9961646 for hepatocytes, and SRR6246979, SRR5714120, and SRR5714119 for macrophages. The human hepatocytes data used were from our single-cell RNAseq results. Data on monocytes were retrieved from SRA under accessions SRR12539860, SRR12539861 and SRR12539862. The Illumina reads were aligned against the NCBI reference genome for mice (GRCm38) or Humans (GRCh38) using HISAT2 v2.1.0[8]. The alignment files were then sorted by SAMtools v.1.6[9] and submitted to HTseq v0.9.1[10] to generate the count file. The count file was then used for the differential expression analysis under R, using DEseq2[7]. The data were normalized and filtered for padj values <0.01.

***Plasmodium* inoculations into mice**

For sporozoite challenge experiments to determine liver-parasite loads, salivary glands of parasitized *A. stephensi* mosquitoes were dissected and sporozoites were isolated, counted and injected 3-5x$10^4$ in 200μL RPMI with 1% mouse serum (Innov-research) retro-orbitally or into the tail vein of mice[11]. For some imaging experiments, parasitized *A. stephensi* mosquitoes (~100/carton) were allowed to bite the abdomen of ketamine anesthetized mice (5/carton) for 6 separate 5-minute intervals.

For infecting of liver-humanized mice, *A. stephensi* infected with the *P. falciparum* NF54 strain were allowed to bite a ketamine-anesthetized liver-humanized mouse for 15 mins. The remaining sporozoites were subsequently isolated from these mosquitoes and inoculated intravenously into the tail vein of the same mouse at 7.5x$10^5$ per mouse. A prophylactic dose of penicillin/streptomycin was administered to these mice to prevent sepsis from the potential injection of non-sterile mosquito-derived material. The animals were euthanized at 30h p.i. to collect the liver.

**Flow Cytometry**

Hepatocyte fractions collected after prefusion, density gradient separation and adhesion to culture plates (dislodged by trypsin, 0.25% trypsin-EDTA, 5 min at 37°C) were stained with anti- CD45 F4/80, CD11c, CD11b or CSF1R (Biolegend) to determine the presence of hematopoietic or Kupffer cells, or to phenotypically characterize them as presented in detail before[1]. Cells were stained for cell surface markers with appropriate antibodies in PBS for 30 minutes prior to washing and resuspending in PBS to analyze by flow cytometry. To determine Caspase-1 activation, *Plasmodium* infected hepatocytes were tested with FAM-FLICA Caspase-1 assay kit (ImmunoChemistry Technologies) as per the manufacturer's protocol. Data were acquired on an LSR Fortessa (BD Biosciences) and analyzed with Flowjo (Treestar).

**Macrophage differentiation**

Bone marrow-derived macrophages (BMDMs) were prepared as described previously[2]. In short, bone marrow cells were grown in L-cell-conditioned IMDM medium (ThermoFisher) supplemented with 10% FCS, 1% non-essential amino acids and 1% penicillin-streptomycin for five days to differentiate into macrophages. On day 5, BMDMs were seeded in 6-well cell culture plates. The next day BMDMs were stimulated with LPS (3.5h) followed by ATP or Nigericin (0.5h), or infected with *St* and the whole-cell lysates or supernatants were collected as indicated.

**Assessment of liver parasite burden**

Liver parasite burden was assessed by quantitative real-time RT-PCR for parasite 18s rRNA in isolated hepatocytes from mice challenged with sporozoites isolated from infected mosquitoes[12]. Total RNA was extracted from isolated hepatocyte fraction at the indicated time points after *Plasmodium* infection, with TRIzol, followed by DNase digestion/cleanup with RNA Clean and Concentrator kit (Zymo Research). 2μg liver RNA per sample was used for qRT-PCR analysis for *Plasmodium* 18S rRNA using TaqMan Fast Virus 1-Step Master Mix (Applied Biosystems). Data were normalized for input to the GAPDH control (hepatocytes) for each sample and are presented as ratios of *Plasmodium* 18s rRNA to GAPDH RNA. The ratios depict relative parasite loads within an experiment and do not represent absolute values.

*Pb-Luc* was used to assess the kinetics of replication and clearance of *Plasmodium* infection in the liver of mice. For bioluminescent detection, mice were injected with D-luciferin (150 mg/kg; PerkinElmer) intraperitoneally and anesthetized using 2% (vol/vol) gaseous isoflurane in oxygen prior to imaging on an IVIS 100 imager (Xenogen). Quantification of bioluminescence and data analysis was performed using Living Image v4.3 software (Xenogen).

**BrdU incorporation**

To generate BrdU incorporated sporozoites, female B6 mice were inoculated with *P*b infected RBCs (1.5-2 x$10^6$/ mouse), i.p., followed by mosquito feeding at days 4 and 5 post-infection. The infected mice were anaesthetized and exposed to cages containing around 200 overnight-fasted female *A. stephensi* mosquitoes. The mosquitoes were then maintained on 12.5% (w/v) sucrose with 1 mg/ml BrdU, at 19-20°C and 80% relative humidity. The sucrose-BrdU solution was made fresh and replaced every 2 days, until the mosquitoes were dissected to obtain the sporozoites on day 21 after the initial feeding. To incorporate BrdU into DNA of B16 melanoma cells, BrdU (10μM) was added to the tissue culture media and cultures maintained for 7 days.

**Cell-death assay**

Lactate dehydrogenase (LDH) release assay to determine cell lysis: Overnight cultures of 5x$10^4$ hepatocytes/ well in 48-well plates were inoculated with 2.5x$10^4$ *P. yoelii* sporozoites. Cell culture supernatants were replaced with fresh media at 4h of incubation after washing the cells twice with media, to remove free sporozoites. Culture supernatant was subsequently collected at various time points and assayed for LDH as a measure of total cytolysis, using the CytoTox 96 Non-Radioactive Cytotoxicity Assay (Promega) according to the manufacturer's instructions. % cytolysis calculated as 100 X (experimental LDH release signal/ maximum LDH release signal).

Propidium Iodide (PI) uptake assay for pyroptosis: PI staining distinguishes programmed cell-death from cell-lysis[13]. To determine the extent of pyroptotic cell death, $3\times10^4$ hepatocytes were seeded per well of an opaque-wall, clear-bottom 96-well plate (Corning). Culture media was replaced with DMEM without phenol red. All culture wells were supplemented with 6 μg/ml PI (Invitrogen) and incubated at 37°C for 40 minutes. Subsequently, the cells were washed thrice with PBS and, the plate was sealed with a clear, adhesive optical plate seal (Applied Biosystems). The frequencies of PI stained cells were determined by flow cytometry or estimated using a plate reader (Sinergy H4, Biotek). % cytolysis calculated as 100 X (experimental group PI signal / maximum PI signal).

**Microscopy**

Liver sections collected from infected ($5x10^5$ *Py* or $7.25x10^5$ *Pf* sporozoites) mice were fixed, permeabilized with 1% Triton X-100 (Fisher Bioscience) and imaged after staining. In addition, cultured hepatocytes in 10u-slide for chemotaxis (ibidi) were imaged live. Images were acquired on SP8 NLO Microscope (Leica) using a 10X/0.40 dry objective (live) or 25x/0.95 water immersion objective with coverslip correction (fixed), as described in detail previously[14]. All images acquired were analyzed using Imaris software (Bitplane).

Caspase-1 (p10, polyclonal, Bioss Antibodies) was used on human hepatocytes and Caspase-1 (p20, Clone Casper-1, Adipogen), GasderminD (Clone EPR19828, AbCam) or AIM2 (Clone EPR18793, AbCam) on mouse hepatocytes. Pfhsp70 (polyclonal, GenWay) was used to mark *Pf.* The following were used to identify *Py*: hep17 (gift from Dr. Scott Lindner, Pennsylvania State University), ACP or CSP (gifts from Dr. Stefan Kappe, Seattle Children's Hospital), along with anti-BrdU (clone BU1/75, ThermoFisher) or CellTrace Violet (CTV, ThermoFisher). For live imaging, hepatocytes were cultured overnight in collagen coated chamber slides (ibidi) and infected with *Plasmodium*. Cultures were maintained in a climate-controlled chamber during imaging. For determining BrdU incorporation in *Plasmodium*, air-dried sporozoites ($10^4$ / well) in "PTFE" printed slides (Electron Microscopy Sciences), or frozen sections from infected livers were used. Samples were fixed with 4% paraformaldehyde/PBS (10 min), permeabilized with Cytofix/Cytoperm buffer (BD Biosciences) for 15 min/ 4°C or 1% tritonX100, washed once with Perm/Wash buffer (BD Biosciences), treated with Cytoperm Permeabilization Buffer Plus (BD Biosciences), for 15 min/ 4°C, washed again with Perm/Wash buffer, treated again with Cytofix/Cytoperm (BD Biosciences) for 15 min/ 4°C. After washing with Perm/Wash buffer again, the samples were treated with DNAse in PBS/BSA for 90 mins at 37°C. The samples were subsequently washed with Perm/Wash buffer and co-incubated with rat monoclonal anti-BrdU antibody (Clone BU-1, Thermofisher) and anti-CSP (sporozoites) or anti-ACP (schizont) antibodies. Subsequently the samples were washed thrice with Perm/Wash buffer, probed with fluorophore conjugated secondary antibodies, washed thrice with Perm/Wash buffer and imaged.

**Inflammasome activity assay**

Caspase-1 activity was assayed in cells as described in detail before[15]. In short, $8x10^5$ hepatocytes from 24h infected or uninfected cultures were lysed and the supernatants were assayed for residual caspase-1 activity by its ability to cleave the fluorogenic substrate Z-YVAD-AFC (Enzo) by incubating at 37°C, 60 min. The fluorescence was evaluated at Ex/Em 400/505 in a microplate reader (SynergyH1, BioTek).

**Transfection of cells**

Hepatocytes ($7\times10^5$ cells) were transfected with various plasmids (at 6 µM concentration) using the Mouse/ Rat hepatocyte Nucleofector kit (Lonza) following the manufacturer's protocol. The transfected cells were transferred to collagen-coated wells,16 hr prior to infection or treatments.
Differentiated BMDMs were detached by treatment with trypsin and resuspended at a concentration of $10^6$ cells per 100 µl or nucleofector solution. These cells were transfected with ON-TARGETplus SMARTpool siRNAs (Dharmacon) using the Mouse Macrophage Nucleofector kit (Amaxa) following the manufacturer's protocol. The transfected cells were transferred to plates and allowed to recover for 24-48h prior to infections or treatments. The DNA extracted from *Plasmodium* or B16 tumor cells lines using Phenol/ chloroform/ isoamyl alcohol precipitation was sonicated at 20% power (Qsonica) with 5s/45s on/off cycle, 8 times, to shear it to uniform 200bp fragments. BMDMs in 6 well plates were transfected using lipofectamine2000 as described before[16].

**Therapeutic regimens**

The following treatment regimens were used in this study: siRNA (corresponding Dharmacon siGENOMESMARTpool): 1nM/mouse, hydrodynamic i.v., -1 or -2dpi, Poly I:C (Invivogen): 100ug/mouse, hydrodynamic i.v., Mammalian expression plasmids: 10µg/ mouse, Anakinra: 10mg/kg, i.v., -1,0,1dpi, Disulfiram (Sigma-Aldrich): 50mg/kg in sesame oil, i.p

**Sequencing**

To determine the sequence of procaspase-1 in hepatocytes, whole hepatocytes from naïve B6 mice were isolated and total RNA prepared as described above; cDNA was synthesized as described in detail before[17] and procaspase-1 gene amplified with the following primers: 5`-atggctgacaagatcctgagggcaaag-3` (F) and 5`-ttaatgtcccgggaagaggtagaaac-3` (R). The amplicon was cloned using TOPO TA cloning kit (Invitrogen) following the manufacturer's protocol and multiple clones sequenced.

**Hydrodynamic delivery of nucleic acids**

Hydrodynamic injections were performed as described in detail before[18]. In short, the desired amount of the siRNA was resuspended in PBS (at 10% volume/ body weight of the mouse) and delivered to the tail vein of mice, using constant pressure, within 7 seconds. The mice were then placed on a warm heating pad and allowed to recover for about 30 mins, before being transferred back into their cages.

**Adeno-associated viral vectors**

Adeno-associated viral vectors were generated by VectorBuilder Inc. by encoding the genes of interest under albumin promoter to limit their expression to hepatocytes[19]. AAV-DJ strain was used as the background for efficient transduction of hepatocytes in vivo[20]. Virus stocks were resuspended in PBS and inoculated intravenously at $1x10^{11}$ GC/ mouse.

**ELISA**

To determine BrdU incorporation in sporozoites, ELISA was performed as described in detail before[21]. In short, $10^4$ sporozoites were plated per well in triplicate, in 96-well flat bottom plates. Air-dried samples were fixed in 4% formaldehyde/PBS (4 min), then in 50/50 methanol/acetone (2 min), washed 3 times in PBS, fixed with Cytoperm Permeabilization Buffer Plus (BD Biosciences), washed 3 times again with PBS and blocked in 1% BSA/PBS (60 min). All wells were incubated with rat monoclonal anti-BrdU antibody (Clone BU-1, ThermoFisher) and DNAse (Sigma) in PBS for 90 min at 37°C, washed three times in PBS, subsequently incubated for 30 min at RT with anti-rat HRP-conjugated secondary antibody (Santa Cruz) diluted 1:5000 in 1% BSA/PBS and again washed three times before developing.

To quantify specific protein levels in culture supernatants, isolated liver or *ex vivo* cultured hepatocytes, ELISA was performed as described in detail before[14]. In short, serial dilutions of supernatants or whole cell lysates (normalized for total protein) were coated in triplicate in 96-well format in 0.1M carbonate bicarbonate buffer (pH 9.6) overnight at 4°C, washed thrice with 0.05%Tween20 in PBS (PBS-T), blocked for 60 min with 1%BSA/PBS, probed with anti-mouse AIM2 (Clone EPR18793, AbCam), anti-Caspase-11 (Clone 17D9, Novus Biologicals) anti-IL-1β (polyclonal, R&D) or anti-IL-18 (Clone 125-25, MBL international) for 60 min at 37°C, washed 5 times with PBS-T, probed with corresponding HRP conjugated secondary antibodies in 1% BSA/PBS and again washed 3 times with PBS-T before developing. All ELISA assays were developed using TMB liquid substrate system (Sigma), stopped with 2N sulfuric acid and then read at 450nm using ELISA microplate reader (Bio-Tek).

##### Western blot and immunoprecipitation assays

Western blots were performed as detailed before[2]. In short, cells were lysed in RIPA Buffer or sample loading buffer containing DTT and SDS. Proteins were run on 12% SDS-polyacrlyamide gel by electrophoresis and transferred to PVDF (Millipore) or Nitrocellulose (BioRad) membranes. After blocking with 5% skimmed milk or Odyssey blocking Buffer (Licor) for 1h at room temperature, the membranes were probed with the primary antibodies: mCaspase-1p20 (Clone Casper-1, Adipogen), mCaspase-1p10 (Clone Casper-2, Adipogen), mIL-1β (Clone D3H1Z, Cell Signaling), mCaspase-11 (Clone 17D9, Cell Signaling), mIL-18 (Clone 39-3F, MBL), mGasdermin D (Clone EPR19828, AbCam), hCaspase-1p20 (Clone D7F10, Cell Signaling), mAIM2 (Clone EPR18793, AbCam) or mASC (F-9, Santa Cruz) at 4°C overnight, washed with tris-buffered saline containing 0.1% Tween-20 (TBST) for 4 times, and incubated at room temperature for 45 min with relevant secondary polyclonal secondary anti-rabbit, anti-rat or anti-mouse chemiluminescent (Jackson Immunoresearch) or IRDye (LICOR) conjugated antibodies. After 4 washes with TBST, the proteins were visualized using a chemiluminescence detection reagent (Millipore) or directly by fluorescence (LICOR).

Immunoprecipitation was carried out as described before[22]. Briefly, the samples along with the relevant antibodies: anti-BrdU (Clone ZBU30, Invitrogen), hCaspase-1 (Clone D7F10, Cell Signaling Technology), mCaspase-1p10 (Clone Casper-2, Adipogen) and protein-G-Dynabeads (Invitrogen) were incubated overnight at 4°C. The cells or culture supernatant samples were immunoprecipitated overnight at 4°C, after which they were washed 3 times with PBS containing 0.1% Tween-20 (PBST). Bound protein was eluted by the addition of 2X SDS-sample buffer, and boiled for 5 min, and then analyzed by Western Blot analysis as described above except that the secondary antibody utilized for immunoprecipitated samples was VeriBlot for IP Detection (Abcam) to get rid of heavy and light chains of the antibody. Mouse or rabbit IgGs (SantaCruz) served as Immunoprecipitation sample controls.

##### Statistical analyses

Data were analyzed using Prism7 software (GraphPad) and as indicated in figure legends.

1 Kurup, S. P. *et al.* Monocyte-Derived CD11c(+) Cells Acquire Plasmodium from Hepatocytes to Prime CD8 T Cell Immunity to Liver-Stage Malaria. *Cell Host Microbe* **25**, 565-577 e566, doi:10.1016/j.chom.2019.02.014 (2019).
2 Gurung, P. *et al.* Chronic TLR Stimulation Controls NLRP3 Inflammasome Activation through IL-10 Mediated Regulation of NLRP3 Expression and Caspase-8 Activation. *Sci Rep* **5**, 14488, doi:10.1038/srep14488 (2015).
3 Roth, A. *et al.* A comprehensive model for assessment of liver stage therapies targeting Plasmodium vivax and Plasmodium falciparum. *Nature Communications* **9**, 1-16, doi:10.1038/s41467-018-04221-9 (2018).
4 Law, C. W., Chen, Y., Shi, W. & Smyth, G. K. voom: Precision weights unlock linear model analysis tools for RNA-seq read counts. *Genome Biol* **15**, R29, doi:10.1186/gb-2014-15-2-r29 (2014).
5 Ritchie, M. E. *et al.* limma powers differential expression analyses for RNA-sequencing and microarray studies. *Nucleic Acids Res* **43**, e47, doi:10.1093/nar/gkv007 (2015).
6 Butler, A., Hoffman, P., Smibert, P., Papalexi, E. & Satija, R. Integrating single-cell transcriptomic data across different conditions, technologies, and species. *Nat Biotechnol* **36**, 411-420, doi:10.1038/nbt.4096 (2018).
7 Love, M. I., Huber, W. & Anders, S. Moderated estimation of fold change and dispersion for RNA-seq data with DESeq2. *Genome Biol* **15**, 550, doi:10.1186/s13059-014-0550-8 (2014).
8 Kim, D., Langmead, B. & Salzberg, S. L. HISAT: a fast spliced aligner with low memory requirements. *Nat Methods* **12**, 357-360, doi:10.1038/nmeth.3317 (2015).
9 Li, H. *et al.* The Sequence Alignment/Map format and SAMtools. *Bioinformatics* **25**, 2078-2079, doi:10.1093/bioinformatics/btp352 (2009).
10 Anders, S., Pyl, P. T. & Huber, W. HTSeq--a Python framework to work with high-throughput sequencing data. *Bioinformatics* **31**, 166-169, doi:10.1093/bioinformatics/btu638 (2015).
11 Kurup, S. P. *et al.* CD11c+ cells acquire Plasmodium from hepatocytes to prime CD8 T cell immunity to liver-stage malaria. *Cell host & microbe* (2019, in press).
12 Arreaza, G., Corredor, V. & Zavala, F. Plasmodium yoelii: quantification of the exoerythrocytic stages based on the use of ribosomal RNA probes. *Experimental parasitology* **72**, 103-105 (1991).
13 DiPeso, L., Ji, D. X., Vance, R. E. & Price, J. V. Cell death and cell lysis are separable events during pyroptosis. *Cell Death Discov* **3**, 17070, doi:10.1038/cddiscovery.2017.70 (2017).
14 Kurup, S. P. *et al.* Regulatory T cells impede acute and long-term immunity to blood-stage malaria through CTLA-4. *Nat Med*, doi:10.1038/nm.4395 (2017).
15 Kummari, E. *et al.* Activity-Based Proteomic Profiling of Deubiquitinating Enzymes in Salmonella-Infected Macrophages Leads to Identification of Putative Function of UCH-L5 in Inflammasome Regulation. *PloS one* **10**, e0135531, doi:10.1371/journal.pone.0135531 (2015).
16 Rathinam, V. A. *et al.* The AIM2 inflammasome is essential for host defense against cytosolic bacteria and DNA viruses. *Nat Immunol* **11**, 395-402, doi:10.1038/ni.1864 (2010).

17 Vijay, R. *et al.* Virus-induced inflammasome activation is suppressed by prostaglandin D2/DP1 signaling. *Proc Natl Acad Sci U S A* **114**, E5444-E5453, doi:10.1073/pnas.1704099114 (2017).

18 Kim, M. J. & Ahituv, N. The hydrodynamic tail vein assay as a tool for the study of liver promoters and enhancers. *Methods Mol Biol* **1015**, 279-289, doi:10.1007/978-1-62703-435-7_18 (2013).

19 Kattenhorn, L. M. *et al.* Adeno-Associated Virus Gene Therapy for Liver Disease. *Hum Gene Ther* **27**, 947-961, doi:10.1089/hum.2016.160 (2016).

20 Grimm, D. *et al.* In vitro and in vivo gene therapy vector evolution via multispecies interbreeding and retargeting of adeno-associated viruses. *J Virol* **82**, 5887-5911, doi:10.1128/JVI.00254-08 (2008).

21 Merrick, C. J. Transfection with thymidine kinase permits bromodeoxyuridine labelling of DNA replication in the human malaria parasite Plasmodium falciparum. *Malar J* **14**, 490, doi:10.1186/s12936-015-1014-7 (2015).

22 Kuriakose, T. *et al.* ZBP1/DAI is an innate sensor of influenza virus triggering the NLRP3 inflammasome and programmed cell death pathways. *Sci Immunol* **1**, doi:10.1126/sciimmunol.aag2045 (2016).

1 Kurup, S. P. *et al.* Monocyte-Derived CD11c(+) Cells Acquire Plasmodium from Hepatocytes to Prime CD8 T Cell Immunity to Liver-Stage Malaria. *Cell Host Microbe* **25**, 565-577 e566, doi:10.1016/j.chom.2019.02.014 (2019).

2 Gurung, P. *et al.* Chronic TLR Stimulation Controls NLRP3 Inflammasome Activation through IL-10 Mediated Regulation of NLRP3 Expression and Caspase-8 Activation. *Sci Rep* **5**, 14488, doi:10.1038/srep14488 (2015).

3 Roth, A. *et al.* A comprehensive model for assessment of liver stage therapies targeting Plasmodium vivax and Plasmodium falciparum. *Nature Communications* **9**, 1-16, doi:10.1038/s41467-018-04221-9 (2018).

4 Law, C. W., Chen, Y., Shi, W. & Smyth, G. K. voom: Precision weights unlock linear model analysis tools for RNA-seq read counts. *Genome Biol* **15**, R29, doi:10.1186/gb-2014-15-2-r29 (2014).

5 Ritchie, M. E. *et al.* limma powers differential expression analyses for RNA-sequencing and microarray studies. *Nucleic Acids Res* **43**, e47, doi:10.1093/nar/gkv007 (2015).

6 Butler, A., Hoffman, P., Smibert, P., Papalexi, E. & Satija, R. Integrating single-cell transcriptomic data across different conditions, technologies, and species. *Nat Biotechnol* **36**, 411-420, doi:10.1038/nbt.4096 (2018).

7 Love, M. I., Huber, W. & Anders, S. Moderated estimation of fold change and dispersion for RNA-seq data with DESeq2. *Genome Biol* **15**, 550, doi:10.1186/s13059-014-0550-8 (2014).

8 Kim, D., Langmead, B. & Salzberg, S. L. HISAT: a fast spliced aligner with low memory requirements. *Nat Methods* **12**, 357-360, doi:10.1038/nmeth.3317 (2015).

9 Li, H. *et al.* The Sequence Alignment/Map format and SAMtools. *Bioinformatics* **25**, 2078-2079, doi:10.1093/bioinformatics/btp352 (2009).

10 Anders, S., Pyl, P. T. & Huber, W. HTSeq--a Python framework to work with high-throughput sequencing data. *Bioinformatics* **31**, 166-169, doi:10.1093/bioinformatics/btu638 (2015).

11 Kurup, S. P. *et al.* CD11c+ cells acquire Plasmodium from hepatocytes to prime CD8 T cell immunity to liver-stage malaria. *Cell host & microbe* (2019, in press).
12 Arreaza, G., Corredor, V. & Zavala, F. Plasmodium yoelii: quantification of the exoerythrocytic stages based on the use of ribosomal RNA probes. *Experimental parasitology* **72**, 103-105 (1991).
13 DiPeso, L., Ji, D. X., Vance, R. E. & Price, J. V. Cell death and cell lysis are separable events during pyroptosis. *Cell Death Discov* **3**, 17070, doi:10.1038/cddiscovery.2017.70 (2017).
14 Kurup, S. P. *et al.* Regulatory T cells impede acute and long-term immunity to blood-stage malaria through CTLA-4. *Nat Med*, doi:10.1038/nm.4395 (2017).
15 Kummari, E. *et al.* Activity-Based Proteomic Profiling of Deubiquitinating Enzymes in Salmonella-Infected Macrophages Leads to Identification of Putative Function of UCH-L5 in Inflammasome Regulation. *PloS one* **10**, e0135531, doi:10.1371/journal.pone.0135531 (2015).
16 Rathinam, V. A. *et al.* The AIM2 inflammasome is essential for host defense against cytosolic bacteria and DNA viruses. *Nat Immunol* **11**, 395-402, doi:10.1038/ni.1864 (2010).
17 Vijay, R. *et al.* Virus-induced inflammasome activation is suppressed by prostaglandin D2/DP1 signaling. *Proc Natl Acad Sci U S A* **114**, E5444-E5453, doi:10.1073/pnas.1704099114 (2017).
18 Kim, M. J. & Ahituv, N. The hydrodynamic tail vein assay as a tool for the study of liver promoters and enhancers. *Methods Mol Biol* **1015**, 279-289, doi:10.1007/978-1-62703-435-7_18 (2013).
19 Kattenhorn, L. M. *et al.* Adeno-Associated Virus Gene Therapy for Liver Disease. *Hum Gene Ther* **27**, 947-961, doi:10.1089/hum.2016.160 (2016).
20 Grimm, D. *et al.* In vitro and in vivo gene therapy vector evolution via multispecies interbreeding and retargeting of adeno-associated viruses. *J Virol* **82**, 5887-5911, doi:10.1128/JVI.00254-08 (2008).
21 Merrick, C. J. Transfection with thymidine kinase permits bromodeoxyuridine labelling of DNA replication in the human malaria parasite Plasmodium falciparum. *Malar J* **14**, 490, doi:10.1186/s12936-015-1014-7 (2015).
22 Kuriakose, T. *et al.* ZBP1/DAI is an innate sensor of influenza virus triggering the NLRP3 inflammasome and programmed cell death pathways. *Sci Immunol* **1**, doi:10.1126/sciimmunol.aag2045 (2016).
